# Membrane Tension Drives Opening of NINJ1 Lesions in Dying Cells

**DOI:** 10.1101/2024.10.29.620849

**Authors:** Ella Hartenian, Elliott M. Bernard, Giulia Ammirati, Hubert B. Leloup, Stefania A. Mari, Morris Degen, Magalie Agustoni, José Carlos Santos, Ivo M. Glück, Gonzalo Cebrero, Mirko Stauffer, Jonne Helenius, Dimitrios Fotiadis, Camilo Perez, Christian Sieben, Virginie Petrilli, Sebastian Hiller, Sylvain Monnier, Kristyna Pluhackova, Daniel J. Müller, Petr Broz

## Abstract

Ninjurin-1 (NINJ1) executes cell lysis by polymerizing into filaments, yet how these form membrane lesions is unknown. Integrating cell-based assays with molecular dynamics simulations, we demonstrate that NINJ1-driven membrane rupture starts with the lateral association of NINJ1 dimers into double filaments. Subsequently, swelling of necrotic cells increases plasma membrane tension, leading to separation of the filament interface and NINJ1 lesion opening. Inhibiting cell swelling prevents this increase in tension, blocking NINJ1-mediated plasma membrane rupture without disrupting NINJ1 oligomerization, implicating membrane tension as key for opening NINJ1 lesions. The NINJ1 homolog NINJ2 oligomerizes but fails to form membrane lesions during cell death due to the greater stability of its dimer. Finally, using atomic force microscopy of NINJ1-containing proteoliposomes, we show that NINJ1 forms large pores by stabilizing membrane edges. In summary, our findings support a model whereby membrane tension drives a zipper-like opening of NINJ1 double filaments, initiating necrotic cell lysis.

## Introduction

A timely, proportionate and correctly localized inflammatory response during tissue damage and infection requires the release of cytokines, chemokines and damage-associated molecular patterns (DAMPs). These recruit and activate immune cells to eliminate pathogens, clear debris and initiate healing^1^. However, aberrant release and sensing of DAMPs can contribute to autoinflammatory disease^2^. An important source of these molecules is dying cells, especially those undergoing necrotic forms of cell death such as necroptosis, pyroptosis and ferroptosis. Apoptosis, classically thought of as an immunologically “silent” form of cell death, can also become a source of DAMPs when the number of apoptotic corpses overwhelms their removal by phagocytes, resulting in secondary necrosis and inflammation^3^.

Pyroptosis and necroptosis are initiated by effector proteins, specifically Gasdermins and mixed lineage kinase domain-like pseudokinase (MLKL) respectively, which form pores in the plasma membrane^4^. While MLKL pores only allow the movement of cations^5^, the Gasdermin D pore, with an inner diameter of approximately 22 nm, permits the secretion of certain cytokines, including IL-1β and IL-18, low molecular weight proteins such as S100A8/S100A9, as well as oxylipins^6–8^. However, the relatively small size of these pores is unable to explain the release of larger DAMPs, including HMGB1, nuclear proteins, and nucleic acids that are associated with necrotic forms of cell death^9^. Thus, for many years, DAMP release was attributed to osmotic pressure buildup, followed by cell swelling and eventual passive plasma membrane rupture (PMR), all of which are hallmarks of cellular necrosis.

In a seminal paper in 2021, Kayagaki *et al.* demonstrated that Ninjurin-1 (NINJ1) is essential for plasma membrane rupture and the release of DAMPs during pyroptosis and secondary necrosis of apoptotic cells^10^. It has subsequently been confirmed that NINJ1 is also important for plasma membrane rupture during a range of cell death modalities, including ferroptosis, cuproptosis, and death induced by pore forming toxins^11,12^. NINJ1 is a 16 kDa plasma membrane protein that is highly conserved in mammals and is widely expressed in various tissues and cell types. At resting state NINJ1 comprises 3 straight transmembrane helices (TM1-3) and forms an autoinhibited, inactive face-to-face homodimer that is stabilized by polar contacts between the amphipathic TM1 helices of each monomer^13^. During cell death the homodimer is thought to dissociate, leading to the formation of NINJ1 filaments of up to several hundred nanometers in length^14–16^. The Cryo-EM structure of active, filamentous NINJ1 is characterized by a distinct kink in TM1, resulting in two separate helices: ɑ1 and ɑ2, of which ɑ1 connects neighboring protomers, thereby stabilizing the filament^14–16^. However, how cell death triggers the dissociation of NINJ1 homodimers, the kinking of TM1, NINJ1 oligomerization, and plasma membrane rupture remains unknown.

The fact that necrotic cells release large DAMPs in a NINJ1-dependent manner suggests that oligomeric NINJ1 causes the formation of large holes in the plasma membrane. How NINJ1 filaments form these lesions has not been experimentally confirmed. Given the amphipathic nature of NINJ1 filaments, several non-mutually exclusive models have been proposed. One possibility is that upon the dissociation of NINJ1 homodimers, NINJ1 monomers associate into single filaments, which form lesions by capping membrane edges with their hydrophobic face and lining the pore channel with their hydrophilic face. This mechanism is reminiscent of the pore formation mechanism proposed for Gasdermin D and other pore-forming proteins^17^. Alternatively, it is possible that NINJ1 forms a double filament in the plasma membrane through association of the hydrophilic faces of single NINJ1 filaments. Subsequent opening of the double filament ‘zipper’ provides a hydrophilic conduit through which DAMPs exit the cell^14^. A third model suggests that circular NINJ1 filaments can solubilize discs of plasma membrane lipids (‘cookie cutter’ model), based on the observation that recombinant NINJ1 filaments can curve and expose the hydrophilic face on their convex side^15,16^. Yet, it remains unclear if any of these models correctly describe the mechanism of NINJ1-induced PMR; and the resulting NINJ1 membrane lesions have yet to be visualized.

In this manuscript, we investigate the mechanism by which oligomeric NINJ1 forms plasma membrane lesions in dying cells. Using ferroptosis as a model to study NINJ1-induced plasma membrane rupture (PMR), we find that necrotic cell swelling is associated with an increase in plasma membrane tension and the gradual opening of NINJ1 lesions. Inhibition of cell swelling using osmolytes blocks the increase in membrane tension, and prevents NINJ1-mediated PMR in ferroptotic, apoptotic, and pyroptotic cells without disrupting NINJ1 oligomerization, thus implicating tension as a key factor controlling the opening of NINJ1 lesions. We next used molecular dynamics (MD) simulations to explore how NINJ1 oligomers form and whether membrane tension can promote their opening. Our simulations show that inactive NINJ1 with a straight TM1 laterally associates into larger oligomers that resemble double filaments. Under tension, these double filaments separate allowing the kinking of TM1 and the transition to the active form of NINJ1, which forms a membrane lesion. Consistent with this model, we observe that NINJ1 mutants with a stabilized dimer interface oligomerize but fail to induce PMR. Furthermore, NINJ2—a close homolog of NINJ1 that is deficient for PMR—oligomerizes but does not induce membrane lesions during cell death, due to the greater stability of its dimeric form. Finally, we use atomic force microscopy (AFM) to visualize NINJ1-induced lesions in liposomal membranes and find that NINJ1 forms large, irregularly shaped pores by stabilizing membrane edges. Together, our findings support a two-step model for NINJ1-driven necrosis, in which NINJ1 oligomerizes to form inactive, membrane-embedded double filaments that act as weak points that open under tension in a zipper-like manner to create large membrane lesions.

## Results

### Cell death-associated swelling coincides with gradual opening of NINJ1 lesions

Since NINJ1 activation occurs during various forms of programmed necrosis, we asked if cell swelling and loss of membrane integrity, two hallmarks of necrosis, are needed for NINJ1 activation. We excluded membrane permeabilization as a signal for NINJ1 activation early in our analysis, as PI uptake, which reports the loss of membrane integrity, required NINJ1 activity in ferroptotic cells ^10,11^ (**Figure S1A**). Instead, we focused on cell swelling as a potential activator of NINJ1. To correlate NINJ1 activation with cell swelling we monitored DRAQ7 uptake into individual cells while using the membrane marker CellBrite to measure cell diameter. Upon induction of ferroptosis, both WT and *Ninj1^−/−^* macrophages swelled with similar kinetics, reaching approximately 160% of their initial diameter (**Figures 1A-D and S1B,C**). Interestingly, we observed that NINJ1 activation, or lesion formation (defined by the sharp increase in DRAQ7 nuclear fluorescence in WT cells), only occurred around 60 minutes after the onset of swelling (**Figure 1B,D; Movie 1**). Cell swelling is driven by ionic imbalance; this observation is thus consistent with previous reports showing that NINJ1 activation in ferroptotic cells is preceded by the formation of small membrane lesions or channels that permit ion fluxes^11^.

**Figure 1.**
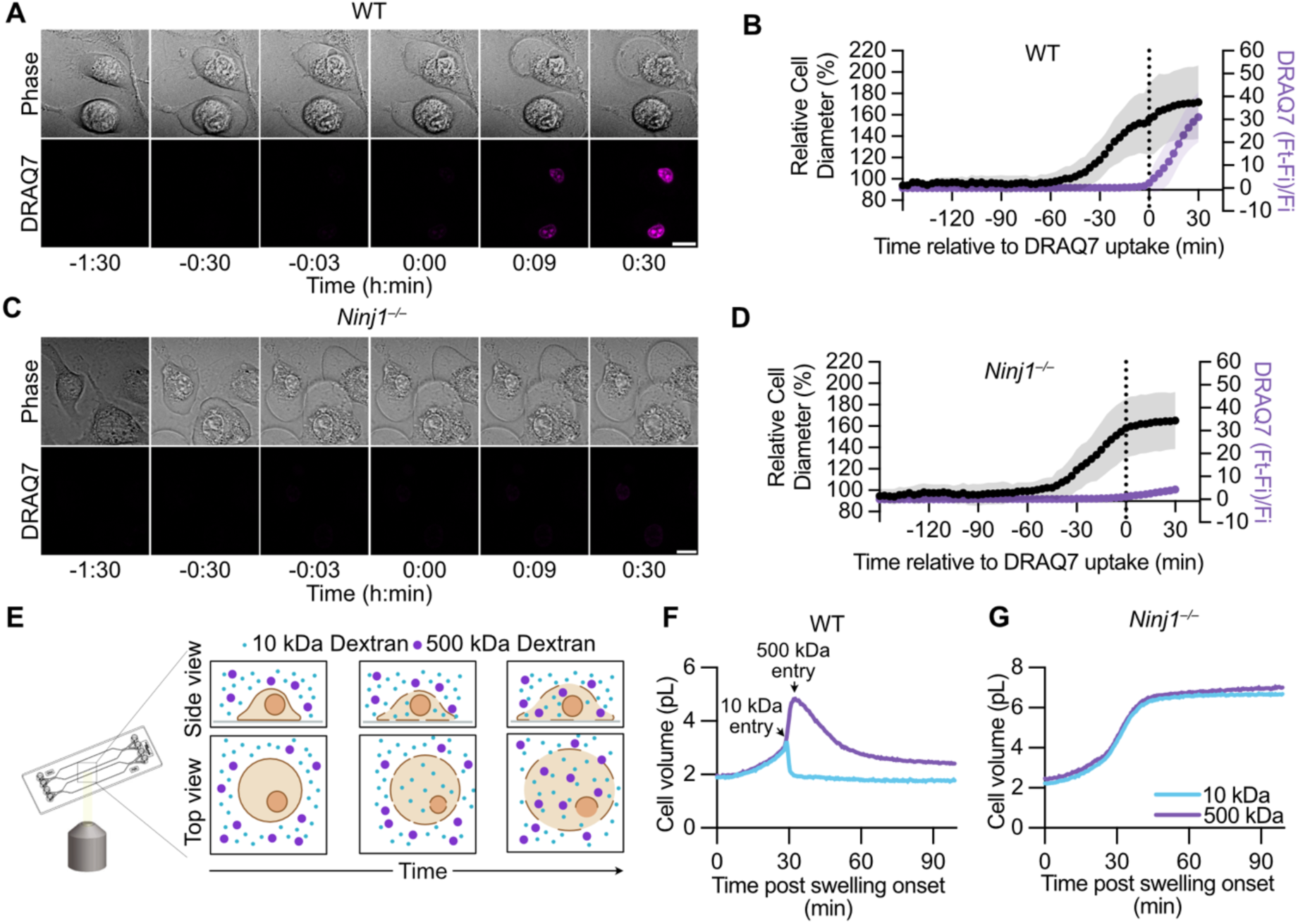
Cell death-associated swelling coincides with gradual opening of NINJ1 lesions. **(A&C)** Stills from live cell imaging of ML162 treated WT (A) and *Ninj1^−/−^* (C) BMDMs in the presence of DRAQ7. Scale bar 10 μm. **(B&D)** Quantification of the diameter of WT (B) and *Ninj1^−/−^*(D) BMDMs treated with ML162 and corresponding DRAQ7 ratios. Time 0 indicates the onset of DRAQ7 uptake. **(E)** Schematic representation of the fluorescence exclusion microscopy (Fxm) approach used to measure cell volume and lesion size. (**F-G)** Cell volume measurements in picolitres (pL) using FXm of WT (F) and *Ninj1^−/−^* (G) iBMDMs treated with cumene hydroperoxide (cumene) in the presence of 10 kDa Alexa647 dextran and 500 kDa FITC dextran. Time 0 indicates the onset of swelling. Images in (A&C) are representative of 3 independent experiments. Data in (B) and (D) are mean ± SD of 20 cells from 3 independent experiments. Data in (F) and (G) are from 1 cell, representative of at least 30 cells measured in 2 independent experiments. For further data see Figure S1.

To more precisely correlate cell volume changes with NINJ1 lesion opening, we utilized a technique based on fluorescence exclusion microscopy (FXm)^18^. Cells are grown in a microfluidic chamber in the presence of small (10 kDa) and large (500 kDa) membrane impermeable fluorescent dextrans, and cell volume is determined by the exclusion of the fluorescent signal (**Figure 1E**). Dye influx, which appears as a sharp drop in the volume trace (dye exclusion is required for volume determination), defines the time point of membrane permeabilization and the size of membrane lesions. FXm measurements demonstrated that, during ferroptosis, the volume of WT immortalized BMDMs (iBMDMs) increased before the entry of the 10 kDa dye, indicative of initial lesion formation, which was then followed minutes later by uptake of the 500 kDa dye (**Figures 1F and S1D,E**). Importantly, between the uptake of the 10 and 500 kD dye there was an additional large increase in cell volume (**Figures 1F (arrows) and S1F,G**). The temporal delay between the uptake of the two dyes could either indicate a gradual growth of NINJ1 lesions by progressive NINJ1 oligomerization, or a gradual opening of NINJ1 lesions, e.g. a transition between a partially- and fully- open state. Notably, we did not observe any uptake of the dyes in *Ninj1^−/−^* iBMDMs and the cells remained swollen until the end of the experiment (**Figures 1G and S1H,I**), corroborating previous findings that NINJ1 executes plasma membrane permeabilization in ferroptotic macrophages^11^. No change in cell volume was detected following treatment with the vehicle control, ethanol (**Figure S1J**). In summary, these results demonstrate that NINJ1 lesions form shortly (within 30-60 min) after cells start to swell. Moreover, the differential entry of small and large dextrans in FXm suggest that NINJ1 membrane lesions vary in size, with the lesion size correlating with the extent of cell swelling.

### Inhibiting cell swelling during cell death prevents the opening of NINJ1 lesions, but not NINJ1 oligomerization

Given the correlation between cell volume and NINJ1 lesion opening, we investigated if swelling is necessary for NINJ1 activation. Experimentally, swelling can be blocked by the addition of hydrophilic molecules (osmolytes), such as large polyethylene glycols (PEGs) or sucrose, to the cell culture medium to reduce the difference in osmolarity between the outside and the inside of the cell. We thus measured cell diameter and DRAQ7 uptake in WT BMDMs undergoing ferroptosis in the presence of large PEGs. In the presence of 10 mM PEG6000 (310 mOsm, compared to 265 mOsm of regular media) the swelling of ferroptotic cells was considerably impaired (**Figures 2A,B; Movie 2**), which was coupled with reduced plasma membrane permeabilization (PI uptake) and rupture (LDH release) (**Figures 2D and S2A**), in agreement with previous findings^12^. Smaller PEGs, as well as sucrose, had a similar effect on swelling, PI uptake and LDH release, even though higher concentrations were necessary to achieve an increase in osmolarity and inhibition of PMR (**Figures 2C,D and S2B,C**). Importantly, while PEGs blocked cell swelling and PMR, they did not prevent cells from dying, as observed by a loss of cellular ATP (**Figure S2D,E**).

**Figure 2.**
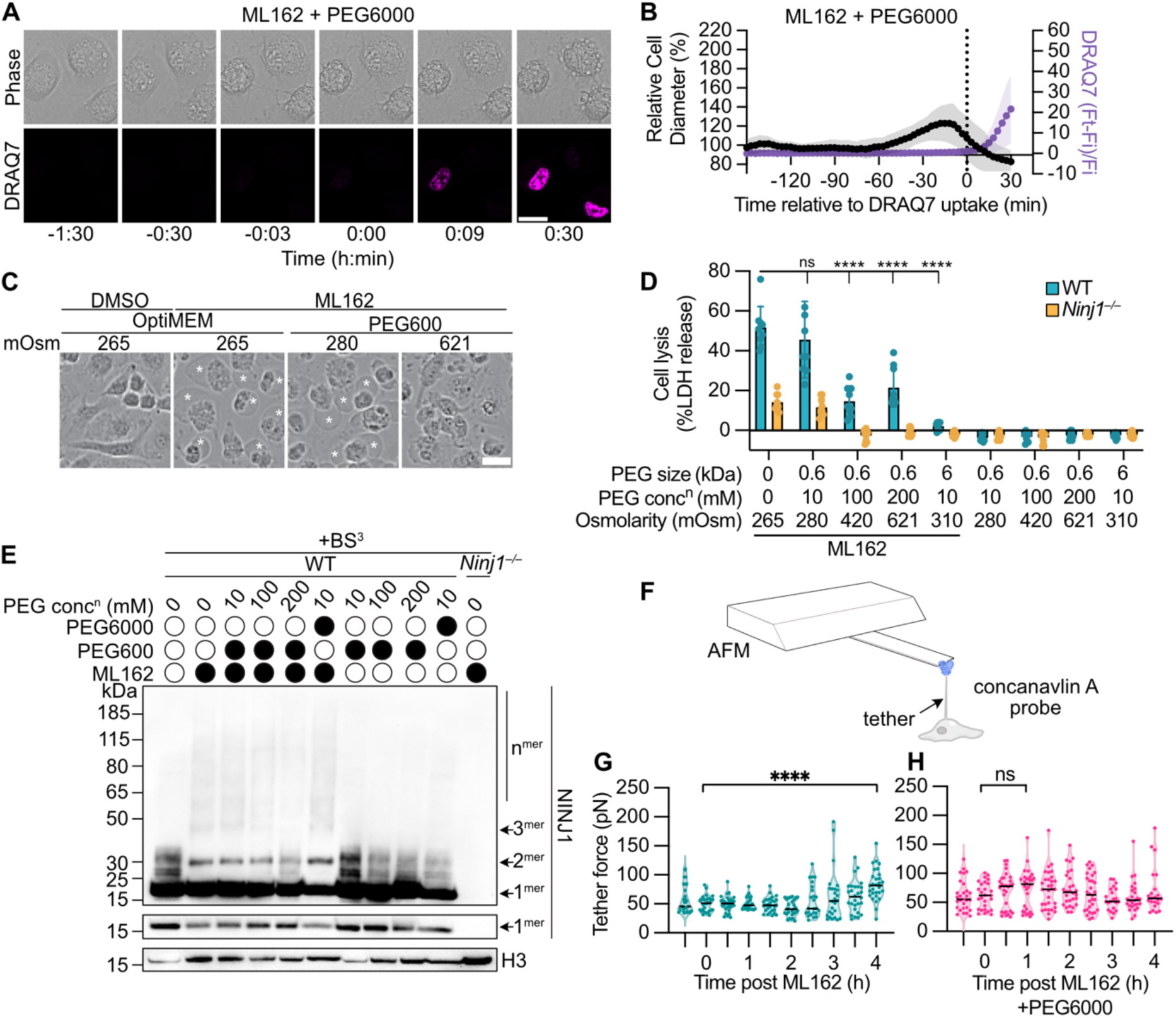
Lesion formation by NINJ1 oligomers requires osmotic swelling. **(A)** Stills from live cell imaging of WT BMDMs treated with ML162 in the presence of 10 mM PEG6000 and DRAQ7. Scale bar 10 μm. **(B)** Quantification of the diameter and corresponding DRAQ7 ratios of WT BMDMs treated with ML162 in the presence of 10 mM PEG6000. Time 0 indicates the onset of DRAQ7 uptake. **(C)** Phase contrast images of WT BMDMs treated or not with ML162 for 4 h in the presence of the indicated osmolarities of PEG600. Asterisks highlight balloons. Scale bar 20 μm. **(D)** Analysis of LDH release from WT and *Ninj1^−/−^* BMDMs treated for 4 h with ML162 in media containing the indicated concentrations of PEG600 or PEG6000. **(E)** Crosslinking immunoblot of WT and *Ninj1^−/−^* BMDMs treated with ML162 in the presence of the indicated concentrations of PEG600 or PEG6000 for 4 h. **(F)** Schematic of the AFM tether pulling technique used to measure membrane tension. **(G-H)** AFM tether force in ML162 treated WT BMDMs in the absence (G) or presence (H) of 10 mM PEG6000. All compounds were added at time 0. Images in (A&C) are representative of 3 independent experiments. Data in (B) are mean ± SD of 20 cells from 3 independent experiments. Data in (D) are mean ± SD from 3 independent experiments, analyzed by 2-way ANOVA followed by Šídák’s multiple comparisons tests comparing all conditions to untreated. Data in (E) are representative of 3 independent experiments. Data in (G-H) are from 3 independent experiments, analyzed by 2-way ANOVA with Dunnett’s multiple comparison test. ns not significant, **** p<0.0001. For further data see Figures S2 and S3.

To further explore how the inhibition of cell swelling prevented NINJ1-dependent PMR, we next tested if PEGs or sucrose abrogated the formation of NINJ1 oligomers, in a manner akin to glycine that blocks LDH release by preventing NINJ1 oligomerization^19^. Unexpectedly, we found that NINJ1 oligomerization during cell death through ferroptosis was largely unaffected by the presence of osmolytes at concentrations preventing LDH release (**Figures 2E and S2F**). Importantly, the tested concentrations of osmolytes alone did not cause any NINJ1 oligomerization, excluding PEGs and sucrose as direct activators of NINJ1 (**Figures 2E and S2F**). Finally, we found that the same osmolytes also inhibited NINJ1-mediated PMR without affecting NINJ1 oligomerization in cells undergoing pyroptosis and secondary necrosis (**Figures S3A-F**).

To test if swelling alone was sufficient to induce NINJ1-dependent PMR, we exposed BMDMs to hypotonic conditions. Although some NINJ1 oligomerization occurred, no NINJ1-dependent LDH release was detected, even at the lowest osmolarity (76 mOsm) (**Figures S3G&H**). Since prolonged hypotonic stress reduced ATP levels in both WT and *Ninj1^−/−^*cells (**Figure S3I**), we concluded that NINJ1 oligomerization was not a direct consequence of swelling but rather caused by the cell death observed under these conditions. Thus, while swelling can ‘functionalize’ or ‘open’ NINJ1 oligomers during cell death, it is not sufficient to induce NINJ1 oligomerization.

### An increase in plasma membrane tension drives NINJ1 lesion opening

A recent study reported that mechanical strain can trigger NINJ1-dependent PMR^9^. Thus, we hypothesized that necrosis-associated swelling and membrane expansion imposes mechanical strain on the plasma membrane, triggering the opening of NINJ1 lesions. Such strain includes lateral membrane tension, which has been shown to increase transiently in response to osmotic swelling of living cells^20^.

To measure tension in the plasma membrane during ferroptosis, we used atomic force microscopy (AFM) with the cantilever functionalized to mechanically pull membrane nanotubes, or tethers, from the plasma membrane (**Figure 2F**). The force required to pull these tethers reports on the tension in the plasma membrane^21^. We observed an increase in the tether force beginning at 3 h post ferroptosis induction, which continued to increase through 4 h in WT BMDMs (**Figure 2G)**. Since tension is related to tether force by a quadratic equation ^22–24^, membrane tension is predicted to increase much more strongly (**Figure S3J,K**). This timeframe correlates with NINJ1 oligomerization and PMR (**Figure S1A, Figure S3L**) and would be consistent with an increase in membrane tension opening closed NINJ1 lesions.

Given that osmolytes, which block cell swelling, prevent the opening of NINJ1 lesions we hypothesized that there would be no increase in membrane tension when osmolytes were present during ferroptosis. Upon ferroptosis induction in the presence of 10 mM PEG6000 the tether forces slightly increased and afterwards returned to untreated values, thus reflecting no increase in membrane tension up to 4 h post ferroptosis induction (**Figure 2H and S3K**).

Overall, these results demonstrate that inhibiting cell swelling prevents NINJ1-mediated PMR without disrupting NINJ1 oligomerization and implicate membrane tension as a key player in the transition of NINJ1 oligomers from an inactive, closed conformation to an active, open state that drives PMR.

### Membrane tension allows the opening of NINJ1 double filaments

NINJ1 lesions have been proposed to form through three mechanisms: (1) single NINJ1 filaments that form pore-like structures (‘pore’ model), (2) single filaments that solubilize membrane discs via their hydrophilic face (‘cookie cutter’ model), or (3) the gradual opening of a double filament formed by two NINJ1 oligomers associated through their hydrophilic face (‘zipper’ model). To investigate how oligomeric NINJ1 forms membrane lesions and how this process is influenced by membrane tension, we employed both coarse-grained (CG) and atomistic molecular dynamics (MD) simulations. CG simulations simplify molecular systems by grouping atoms into “beads on a string,” while atomistic simulations calculate the motions and interactions of individual atoms in detail^25,26^.

We first modelled a single filament comprised of 20 NINJ1 proteins in their four-helix, TM1-kinked, active conformation (PDB:8SZA^15^) and observed that such filaments spontaneously permeabilize membranes. This started with the formation of small lesions caused by the receding of phospholipids from the hydrophilic face of the filament (**Figures 3A and S4A**). These small lesions rapidly expanded, causing the filament to bend and the ends to connect, forming a pore-like structure stabilized by a single filament of NINJ1. To test if membrane tension affected this process, we repeated the simulations of these 20mer NINJ1 single filaments under 5 mN/m of lateral tension. While higher than the tension in biological membranes, this represents a magnitude commonly employed in MD simulations to study the effects of tension on membranes and proteins in the short timescales compatible with computing availability^27,28^. Applying this membrane tension did not lead to observable differences in the formation of pores by a single NINJ1 filament (**Figures 3A and S4A**). Notably, in none of the thirteen single filament simulations did NINJ1 filaments bend around their hydrophobic side to form rings around lipids, as would be expected in a cookie cutter model. Since we observe a tension dependence in experiments (**Figure 2**), a mechanism of pore formation by a single NINJ1 filament would be inconsistent with the experimental data.

**Figure 3.**
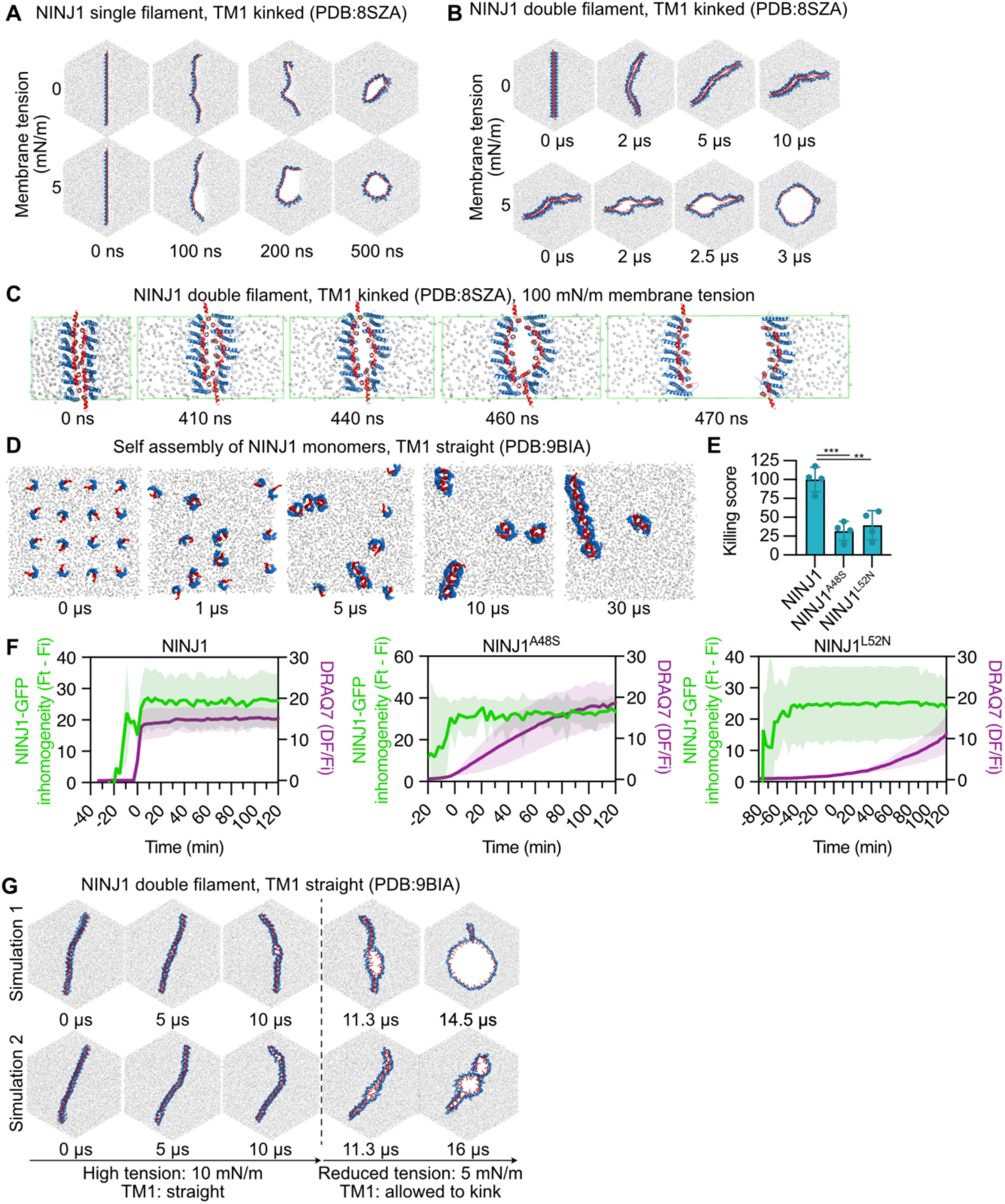
Membrane tension allows the opening of NINJ1 double filaments. **(A)** Snapshots from CG MD simulation of a 20mer NINJ1 single filament (PDB:8SZA) in a membrane in the absence (n=8) or presence (n=5) of 5 mN/m membrane tension. Individual hexagons show snapshots at the indicated time points from the simulations. **(B)** Snapshots from CG MD simulation of a 20mer NINJ1 double filament (PDB:8SZA) arranged with their hydrophilic faces towards each other in the absence (n=3) or presence (n=3) of 5 mN/m membrane tension. **(C)** Snapshots from an all-atom MD simulation of an endless periodic NINJ1 (PDB: 8SZA) double filament with 100 mN/m membrane tension (n=30). **(D)** Snapshots from CG MD simulation of monomeric NINJ1 (PDB: 9BIA) with snapshots at the indicated times of the simulation (n=3). **(E)** Killing score of human (h)NINJ1 and the indicated mutants upon overexpression in HEK293T cells. **(F)** Inhomogeneity and DRAQ7 ratio analysis of cumene treated HeLa cells expressing GFP-tagged hNINJ1 WT and the indicated mutants. **(G)** Snapshots from two CG MD simulations of a 20mer NINJ1 (PDB:9BIA) double filament with a straight TM1 for 10 µs in the presence of 10 mN/m membrane tension, followed by a change in the simulation parameters with a reduction in tension to 5 mN/m, allowing TM1 to kink for the remainder of the simulation. (n=3). In all MD simulations red represents TM1 and blue the remaining structured part of the protein. The lipid phosphates are shown as white spheres. For further data see Figure S4. Data from E are from three independent experimentas and are analyzed with a one-way Anova with Dunnett’s multiple comparison test. *** p=0.0006, ** p=0.0018. Data from F-G are from three independent experiments with 10-15 cells analyzed per condition.

We next simulated a NINJ1 double filament, composed of two 20mer filaments of TM1-kinked NINJ1 associated via their hydrophilic faces, as postulated in the zipper model (**Figures 3B and S4B**). In the absence of membrane tension, the double filament exhibited minor dynamic fluctuations at the filament-filament interface, but never a complete separation of the two filaments (**Figures 3B and S4B; Movie 3**). Strikingly, when membrane tension was applied the pore behavior of the double filament completely changed. Small separations at the filament-filament interface quickly widened and resulted in the complete separation of the two filaments to form a stable pore (**Figures 3B and S4B; Movie 4**). To verify the opening of the double filament, we simulated an endless periodic NINJ1 double filament with kinked TM1 in all-atom mode, which describes the intermolecular more precisely. These simulations came to the result that NINJ1 double filaments open specifically in response to membrane tension (**Figures 3C and S4C,D**). The MD simulations of a NINJ1 double filament under membrane tension are thus fully consistent with our observations in cells, where osmolytes that block increases in membrane tension also prevent the transition of NINJ1 oligomers to an open conformation. Collectively, our data suggest that NINJ1 lesions most likely arise from the tension-dependent opening of NINJ1 double filaments, and not from a single NINJ1 filament.

### Assembly of double filament NINJ1 oligomers occurs independently of dimer dissociation

We next asked how a NINJ1 double filament forms. We speculated that a lateral association of autoinhibited NINJ1 dimers could give rise to an oligomer resembling a closed double filament in which TM1 is straight. Indeed, AlphaFold2 predicts a double filament NINJ1 oligomer with straight TM1 and the hydrophilic face of each filament forming the interface^14^. Furthermore, tetramers and some larger oligomeric forms of NINJ1 are readily observed in live cells, indicating that some dimer-dimer interactions occur at steady state^10,13^. CG simulations starting with wild-type monomeric, TM1-straight NINJ1 (WT model based on NINJ1 K45Q (PDB: 9BIA^13^)), showed that such closed double filaments indeed form spontaneously: NINJ1 monomers assemble into dimers, tetramers, hexamers, octamers, and even 12mers (**Figure 3D; Movie 5**). Moreover, simulations initiated from NINJ1 tetramers, i.e. laterally associated NINJ1 dimers, showed self-assembly into 16mers (**Figure S4E**), further indicating that autoinhibited dimeric NINJ1 can spontaneously assemble larger oligomers.

We thus hypothesized that previously reported mutations that stabilize face-to-face interactions of NINJ1 dimers would reduce cytotoxicity without affecting NINJ1 oligomerization. We selected two such mutants^13^, NINJ1^A48S^ and NINJ1^L52N^, and confirmed that, unlike WT NINJ1, they did not induce spontaneous cell lysis upon overexpression in 293T cells (**Figure 3E**). To measure oligomerization of these mutants we expressed them with a GFP tag in HeLa cells, induced ferroptosis and monitored GFP signal inhomogeneity, as well as DRAQ7 uptake, a readout of PMR (**Figure 3F**). While the expression of NINJ1^A48S^ or NINJ1^L52N^ resulted in slower DRAQ7 uptake when compared to cells expressing WT NINJ1, we observed a similar increase in GFP inhomogeneity for all three proteins, indicating that autoinhibited mutants of NINJ1 formed oligomers upon ferroptosis induction (**Figure 3F**). Taken together, these results demonstrate that the dissociation of NINJ1 dimers is not required for the oligomerization of NINJ1 during cell death. Instead, NINJ1 oligomers are formed through the lateral association of homodimers. Since separation of NINJ1 dimers is required for PMR^13^, this finding also implies that homodimer separation must occur after oligomers have formed.

We next tested if membrane tension could separate NINJ1 double filaments with a straight TM1, and if TM1 could kink after filament separation. In the absence of membrane tension, double filaments formed by 2×20mers of NINJ1 with a straight TM1 were stable over the course of 10 µs of CG simulations (**Figure S4F**). Even with 10 mN/m of membrane tension – double what was required to separate double filaments with kinked TM1 – only small local openings between the individual filaments appeared, which then resealed **(Figure 3G, Movie 6**). This indicates that the TM1-straight double filament is more stable than the TM1-kinked double filament. We thus speculated that the kinking of TM1 could weaken the polar interactions between opposing dimers within the filament to facilitate zipper-like opening and therefore continued the simulation allowing TM1 to kink and simultaneously reducing the membrane tension to 5 mN/m (**Figure 3G, Movie 6**). Under these conditions we observed that the formation of small local openings allowed TM1 conformational flexibility, weakening inter-filament contacts and allowing membrane tension to fully separate the double filament and form a pore. All-atom simulations of periodic endless NINJ1 double filaments initiated with a straight TM1 demonstrated that the application of membrane strain caused the intracellular side of the filaments to partially separate, whereas the extracellular side remained closely associated (**Figure S4G**). Quantifying the distance between TM3 and TM4 of opposing protomers at the intracellular side revealed an average distance of 2.0 nm in the absence of tension, increasing to 2.6 nm once tension was applied (**Figure S4H**). Thus, membrane strain partially opens the intracellular side of inactive NINJ1 double filaments, generating space that would be needed for TM1 kinking and eventual opening. Notably, due to the limited length of our simulations (4-5 µs at all-atom resolution) the straight TM1 of NINJ1 did not fully transition into its kinked, protomer-bridging state, which would lead to further separation of the filaments and their opening (**Figure S4G,H**).

In summary, these results support a model where NINJ1 forms membrane lesions in dying cells through a stepwise process. First, autoinhibited NINJ1 dimers associate laterally into a double filament oligomer. Next, membrane tension separates NINJ1 dimers within this double filament, initially generating small openings that allow straight TM1 to kink, thus weakening interactions between NINJ1 dimers and further promoting the separation of the double filament. At the same time, kinked TM1 stabilizes the interactions between the neighboring NINJ1 protomers, resulting in a filament strong enough to stabilize membrane edges to form a lesion.

### Stability of NINJ2 dimers governs their inability to induce PMR

Since our model suggests that the stability of NINJ1 dimers determines whether NINJ1 oligomers can permeabilize membranes, we next examined NINJ2, which shares 67% sequence identity with NINJ1 (**Figure S5A**). As recombinant NINJ2 forms filaments with a structure similar to NINJ1 *in vitro* but NINJ2 does not cause cell lysis^10,15^, we hypothesized that NINJ2 forms oligomers that remain in a closed state during cell death. In support of this, we observed that NINJ2-GFP transitioned from a diffuse membrane localization to a punctate pattern in HeLa cells undergoing ferroptosis, indicative of oligomerization (**Figure 4A**). We quantified the size distribution of NINJ2 clusters (radius of gyration (Rg)) by total internal reflection fluorescence (TIRF) stochastic optical resolution microscopy (STORM), which confirmed that NINJ2 clusters in ferroptotic cells were significantly larger than NINJ2 structures found in the untreated control cells (**Figures 4B and S5B**). We next compared NINJ1 and NINJ2 oligomerization kinetics using inhomogeneity analysis as done previously^14^ and found that both proteins oligomerized with comparable speed (**Figure 4C**). However, while NINJ1 induced rapid DRAQ7 uptake, NINJ2-GFP expressing cells acquired DRAQ7 slowly, similar to cells expressing the control protein HA^TMD^-GFP^14^ (**Figure 4D**), indicating that the observed DRAQ7 uptake is largely NINJ2-independent.

**Figure 4.**
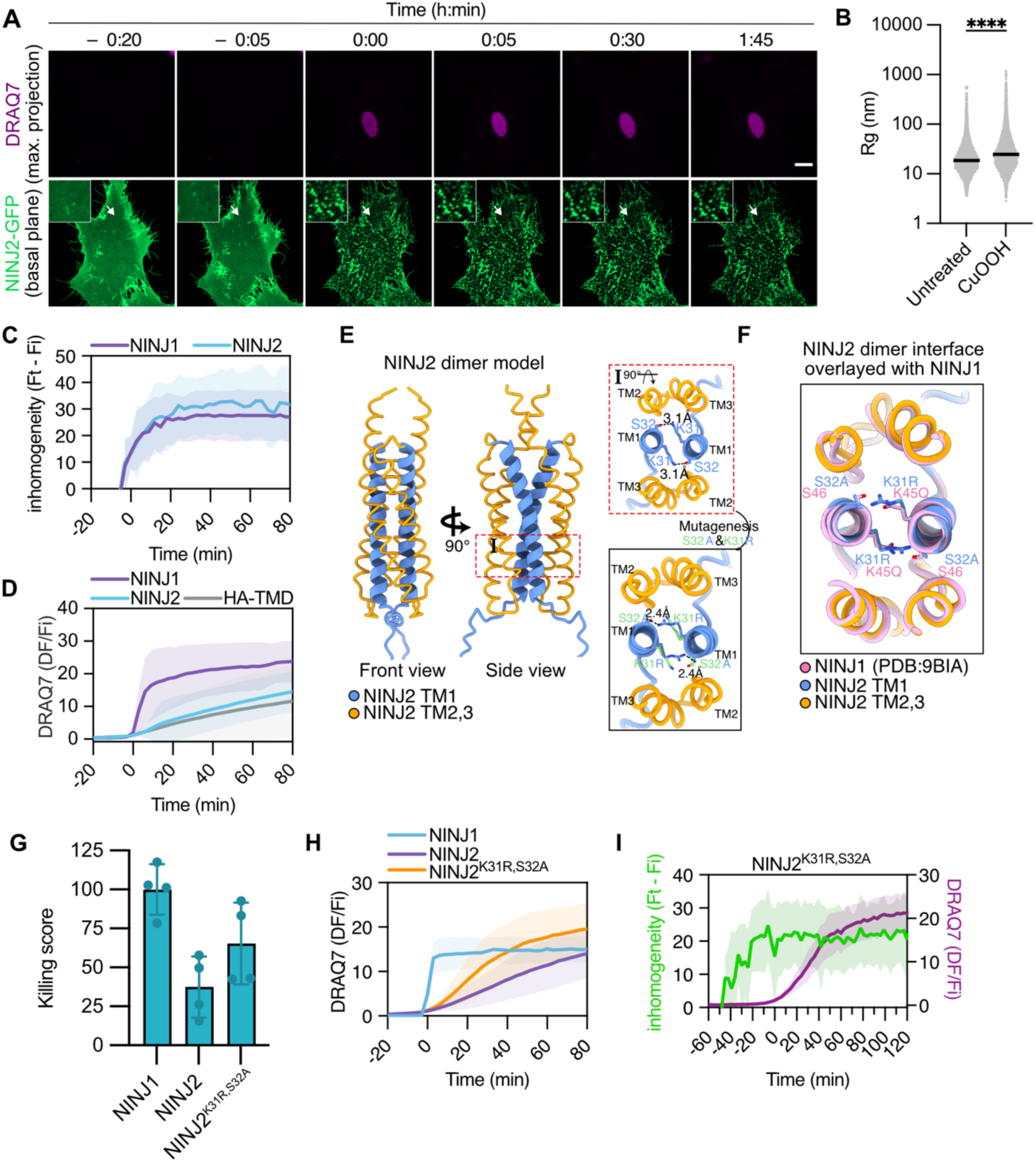
Stability of the NINJ2 dimer determines its inability to induce PMR. **(A)** Stills from live cell imaging of *NINJ1/2* dKO HeLa cells expressing NINJ2-GFP after treatment with cumene in the presence of DRAQ7. Images show NINJ2-GFP expression at the basal plane and the influx of DRAQ7 with a maximum intensity projection. Time 0 indicates the onset of DRAQ7 uptake. Scale bar 10 μm. **(B)** Radius of gyration (*R*g) for each identified NINJ2 cluster from STORM images of untreated or cumene treated HeLa cells expressing NINJ2-GFP. Plots show the distribution of all identified clusters from two independent experiments; lines indicate median values. **(C)** Quantification of the inhomogeneity distribution of NINJ1- or NINJ2-GFP in cumene treated *NINJ1/2* dKO HeLa cells where time 0 indicates the half maximum inhomogeneity of each protein. **(D)** DRAQ7 ratio measured over time of the indicated constructs from live cell imaging performed on cumene treated *NINJ1/2* dKO HeLa cells. Time 0 indicates the onset of DRAQ7 uptake. **(E)** AlphaFold3 model of the NINJ2 dimer interface using the structure of NINJ1 (PDB: 9BIA) for mapping. TM1 is modeled in blue and TM2 and TM3 are modeled in yellow. The interaction of residues K31 and S32 at the protomer interface are shown in the red box, with the modeled interaction of the mutants K31R and S32A shown in the black box. Distances between key residues are indicated. **(F)** The NINJ2 dimer interface, overlaid with the structure of the autoinhibited dimer structure of NINJ1 K45Q (PDB: 9BIA, pink). **(G)** Killing score of human NINJ1, NINJ2 and NINJ2^K31R,S32A^ upon overexpression in HEK293T cells. **(H)** DRAQ7 ratio measured over time of the indicated constructs from live cell imaging performed on cumene treated HeLa cells. Time 0 indicates the onset of DRAQ7 uptake. **(I)** Quantification of the inhomogeneity distribution of NINJ2^K31R,S32A^-GFP in cumene treated *NINJ1/2* dKO HeLa cells and the influx of DRAQ7 (purple). (A, C, D, I & J) are from 3 independent experiments with 16-30 cells each and (C, D, I & J) show the mean ± SD. H is mean ± SD from 4 independent experiments. For further data see Figure S5.

We therefore hypothesized that during cell death NINJ2 assembles into higher-order oligomers that remain in an inactive, or “closed,” conformation, consistent with Sahoo et al’s observation that purified NINJ2 oligomers are defective for lesion formation^15^. For NINJ2 oligomerization to occur through a zipper-like mechanism, NINJ2 must form face-to-face homodimers, as was reported for NINJ1^13^. Indeed, crosslinking analysis demonstrated that endogenous NINJ2 in MOLT-4 cells forms dimers under resting conditions (**Figure S5C**). To see if we could destabilize the NINJ2 dimer interface and promote PMR, we modeled an inactive NINJ2 dimer based on the autoinhibited NINJ1 structure (**Figure 4E**) and identified residues K31 and S32 as candidates whose mutation could destabilize the NINJ2 dimer, analogous to the function of homologous residues in autoinhibited NINJ1 (**Figure 4F**). Consistent with this hypothesis, we found that expression of NINJ2^K31R,S32A^ caused plasma membrane rupture to higher levels than WT NINJ2, despite both proteins being expressed to similar levels (**Figures 4G and S5D**). Moreover, both NINJ1 and NINJ2^K31R,S32A^ caused DRAQ7 uptake in HeLa cells during ferroptosis, while NINJ2^WT^ failed to do so (**Figure 4H**). Importantly, all three proteins – NINJ1, NINJ2, and NINJ2^K31R,S32A^ – oligomerized with similar kinetics (**Figure 4C, I**). LDH release mediated by NINJ2^K31RS32A^ did not attain levels reached by NINJ1, implying that other regions of NINJ2, either within the structured part or unstructured N-terminus, may be important for the increased stability of the double filament interface. NINJ2 chimeras carrying portions of the extended N-terminal region of NINJ1 did not facilitate DRAQ7 uptake, indicating that the unstructured NT is not a determinant of dimer stability (**Figure S5E-F**). In summary, we show that NINJ2 forms higher-order oligomers in dying cells, similar to NINJ1, but these remain inactive due to the greater stability of the NINJ2-NINJ2 filament interface.

### NINJ1 oligomers stabilize membrane edges to form irregularly shaped lesions in biological membranes

Our MD simulations suggest that NINJ1 remains inserted in the plasma membrane after lesions have formed, which contradicts reports that NINJ1 liberates membrane discs to form lesions (i.e. the ‘cookie cutter’ model)^15,16^. To validate these findings experimentally, we asked if NINJ1 oligomers are released from dying cells. Crosslinking immunoblots showed that upon induction of pyroptosis, ferroptosis or apoptosis the majority of NINJ1 dimers, trimers and larger oligomers remained associated with the cell pellet (**Figure 5A**). Supernatant fractions contained mainly monomeric NINJ1, or small amounts of oligomers in the case of apoptosis, but these were negligible compared to the amount of oligomeric NINJ1 in the cell pellet. David *et al.* also reported the release of large rings of NINJ1-GFP from dying cells, which were interpreted as membrane discs released by oligomeric NINJ1^16^. However, the large size of these rings (500–1000 nm) contrasts sharply with the much smaller, heterogeneous NINJ1-lipid structures formed upon incubation with liposomal membranes, which are no more than 50 nm in size^15,16^. Speculating that NINJ1 rings represent exosomes or vesicles, which are commonly released from dying cells, we imaged HeLa cells expressing NINJ1-mCherry and the control protein HA^TMD^-GFP. Consistent with David *et al*., we observed rings of NINJ1-mCherry, but in all cases the NINJ1 signal co-localized with HA^TMD^, suggesting that these were not lipid discs, but indeed vesicle-like structures (**Figure 5B**), which was confirmed by Z-stack analysis (**Figure S6A**).

**Figure 5.**
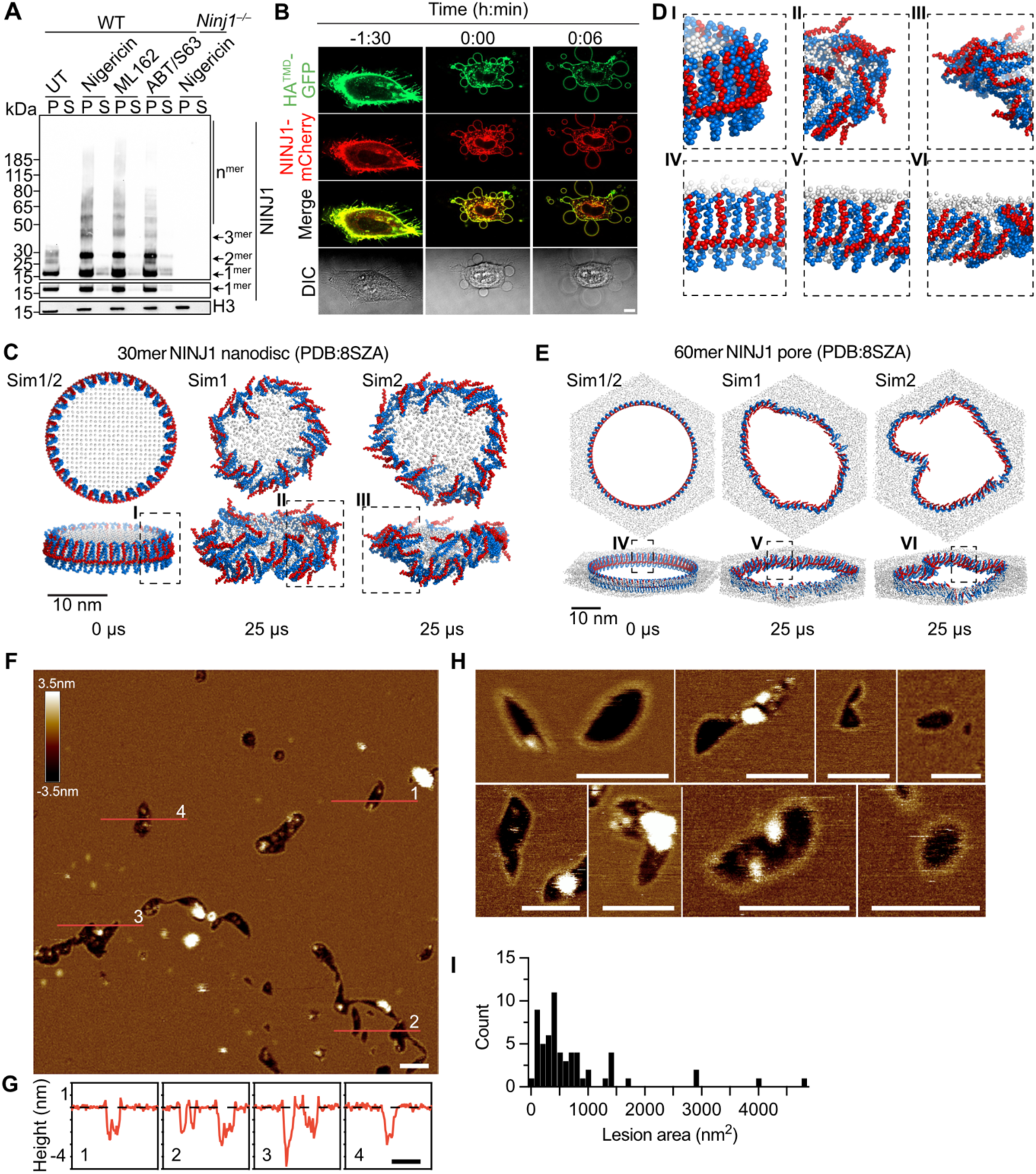
NINJ1 oligomers stabilize membrane edges to form irregular shaped lesions in biological membranes. **(A)** Crosslinking immunoblot of NINJ1 oligomerization in WT and *Ninj1^−/−^*BMDMs left untreated or treated with nigericin (pyroptosis, 2 h), ML162 (ferroptosis, 4 h) or ABT737 + S63845 (apoptosis, 6 h). Cell pellet (P) and supernatants (S) were processed separately. 15% of the cell pellet and 100% of the supernatant sample was loaded and all samples are treated with the crosslinker BS^3^. **(B)** Stills from live cell imaging of cumene treated HeLa cells expressing HA^TMD^-GFP and NINJ1-mCherry in the presence of DRAQ7. Time 0 indicates the onset of DRAQ7 uptake. Scale bar 10 μm. **(C)** CG MD simulations of lipid nano disc surrounded by a 30mer NINJ1 single filament (PDB:8SZA). 25 μs timepoints from two different simulations are shown (n=6). **(D)** Closeups showing the structure of the NINJ1 filament in the corresponding indicated regions of the simulations in C & E. **(E)** CG MD simulations of a pore formed by a 60mer NINJ1 single filament (PDB:8SZA). 25 μs timepoints from two different simulations are shown (n=5). Red represents TM1 and blue the remaining structured part of the protein. The lipid phosphates are shown as white spheres. **(F)** AFM topograph of NINJ1 proteoliposomes adsorbed on freshly cleaved mica. Scale bar 50 nm **(G)** Height profiles of the membrane lesions corresponding to the red lines from (F). Black dashed lines indicate the SLM surface (0 nm height). The full-range color scale of the topograph corresponds to a vertical scale of 6 nm. Scale bar, 50 nm. **(H)** Gallery of AFM topographs showing NINJ1 lesions in lipid membranes, prepared and analyzed as in (F). **(I)** Population analysis showing the size distribution of NINJ1 lesions in liposomal membranes. Data in A is representative of 3 independent experiments. Data in B is representative of 2 independent experiments. Data in F-G is representative of 12 experiments. For further data see Figure S6.

To further compare the pore and cookie cutter models, we used CG MD to simulate the two systems: a lipid nanodisc surrounded by a 30mer TM1-kinked NINJ1 filament, and a NINJ1 pore composed of a 60mer TM1-kinked filament. In the nanodisc simulations, pronounced curvature toward the filament’s hydrophobic face destabilized side-to-side interactions between adjacent NINJ1 subunits, ultimately leading to the disruption of the nanodisc architecture (**Figure 5C**). During the MD simulations, NINJ1 remained attached to the nanodisc, however many holes in the filament structure appeared, with NINJ1 monomers translocating onto the nanodisc surface on both sides (**Figures 5D, panel I-III and S6B**).

In contrast, the pore simulations in which NINJ1 filaments feature a curvature towards the hydrophilic face maintained the filament structure much better (**Figures 5E and S6C**). As observed previously^14^, NINJ1 pores were highly dynamic, while stably maintaining pores with diverse shapes that included regions with both convex and concave curvature. In the convex regions, where the filament bent toward its hydrophobic face, we frequently observed local disruption of the filament structure, further supporting the notion that such curvature is poorly tolerated by filamentous NINJ1.

To provide experimental insights into the mechanism of NINJ1 lesion formation, we next visualized NINJ1 lesions in model biological membranes. We previously showed that recombinant NINJ1 permeabilizes liposomal membranes^14^, thus we examined the NINJ1 lesions formed in liposomes using force distance-based atomic force microscopy (FD-based AFM) – a high-resolution imaging technique we previously used to determine the mechanism of gasdermin pore formation^29,30^. We incubated liposomes composed of 1-palmitoyl-2-oleoyl-sn-glycero-3-phosphoethanolamine (POPE) and 1-palmitoyl-2-oleoyl-sn-glycero-3-phospho-(1’-rac-glycerol) (POPG) lipids with recombinant NINJ1 and adsorbed the sample onto freshly cleaved mica. The AFM topographs, recorded in buffer solution, showed that, upon adsorption to mica, the liposomes opened and formed single-layered membrane patches. At higher resolution, the topographs showed various irregular membrane lesions, most framed by protein borders, although some lesions lacking a protein border were also observed (**Figure 5F**). No lesions could be observed in control samples where NINJ1 was not added (**Figure S6D**). Closer examination revealed that NINJ1 protrudes 0.56 ± 0.11 nm (mean ± SD, *n* = 80) from the lipid surface (**Figure 5G**), forming an almost continuous border. The depth of the lesions was 4.2 ± 0.50 nm (mean ± SD, *n* = 47), which is sufficient to completely penetrate the lipid membrane. To gain further insight into the organization of the proteins bordering the lesion, we measured the distance between individual subunits and found that these were spaced 2.8 ± 0.5 nm (mean ± SD, *n*= 25) apart, which is somewhat larger than the spacing based on the structure of the active TM1-kinked NINJ1 filament solved in the absence of a membrane (≈ 2.1 nm in the CryoEM structure, PDB: 8SZA). Measuring the area of NINJ1-induced lesions showed high variability, ranging from 40 to 4800 nm^2^, with an average size of 735.9 nm^2^ (**Figure 5I**). Thus, NINJ1-induced lesions in liposomes are, on average, twice as large as Gasdermin D pores (inner diameter 21 nm, ≈ area 350 nm^2^)^6^, however due to their heterogeneity some can be one order of magnitude larger. Given the average distance between NINJ1 subunits, our data suggest that the filaments forming these lesions range from 12 to 140 subunits, consistent with the size of filaments modeled in our MD simulations. Taken together, these data are consistent with NINJ1 oligomers remaining associated with biological membranes once NINJ1 lesions have formed. The borderless lesions could arise either from the absence of protein in these few examples, or different conformations or flexibility of NINJ1 that could prevent them from being imaged by AFM. Nevertheless, the large size of some NINJ1 lesions support previous observations that NINJ1 is needed to release large DAMPs that cannot pass through other membrane pores.

## Discussion

Since the discovery of NINJ1, how it ruptures the plasma membrane of dying cells has been a subject of great interest, with multiple triggers for oligomerization and mechanisms of lesion formation proposed ^14–16,31^. Here, combining cell biology, biophysical and *in silico* experiments we propose a comprehensive mechanism for NINJ1-induced PMR (**Figure 6**). Firstly, using cell biology and MD simulations we find that autoinhibited NINJ1 dimers associate laterally leading to the formation of inactive NINJ1 double filaments, with the hydrophilic face sequestered at the interface. Subsequently, using microscopy and biophysical measurements of the cell membrane we see that cells swell and tension in the plasma membrane increases, leading to the separation of the double filament and kinking of TM1, stabilizing the interaction between neighboring NINJ1 protomers^14–16^. MD simulations show that the opening of NINJ1 double filaments results in pore-like membrane lesions that are bordered by a NINJ1 filament. High-resolution AFM identifies NINJ1 framing large lesions in liposomal membranes, which would allow DAMP release and recognition of dead cells by dendritic cells for antigen presentation^10,32^.

**Figure 6.**
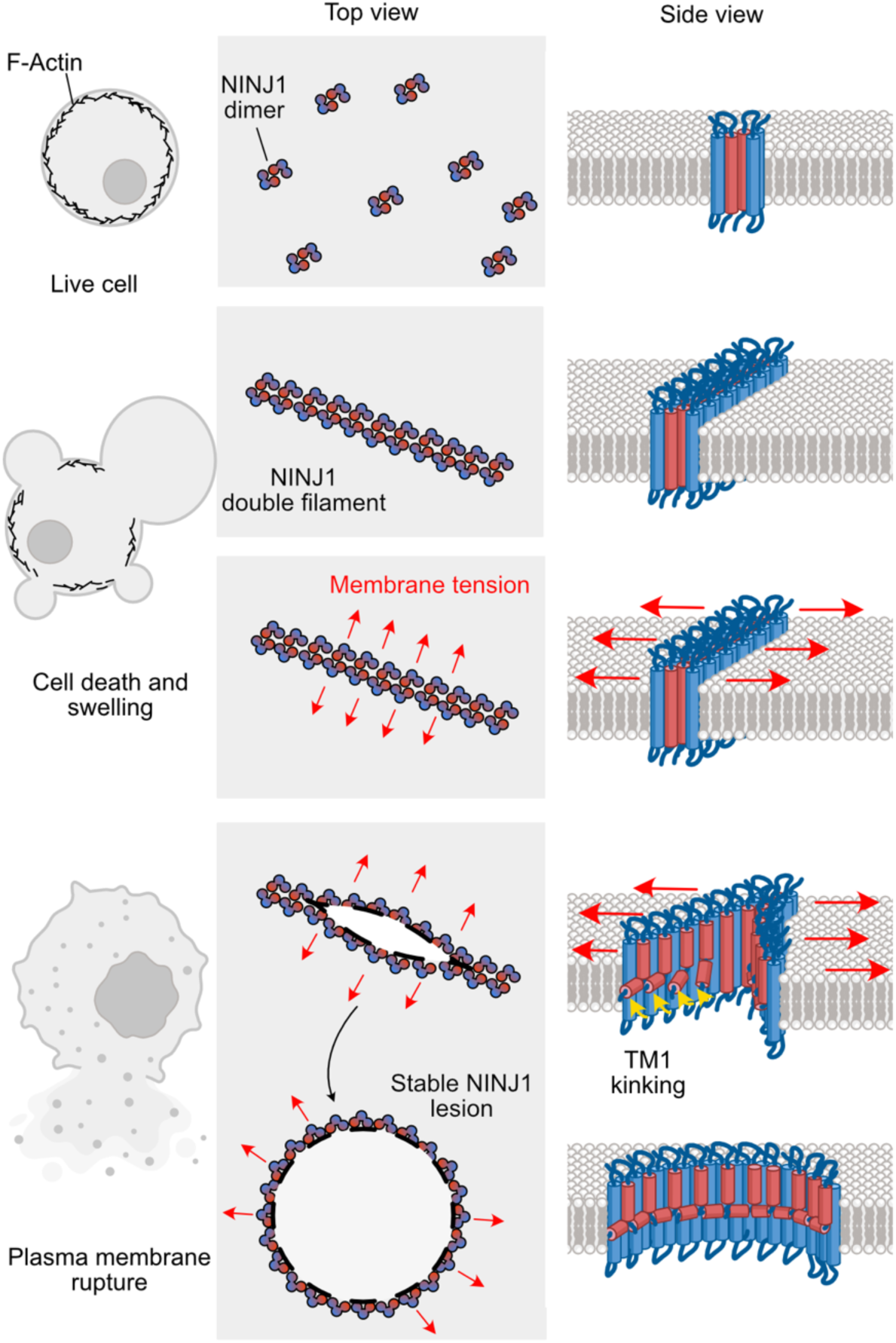
Model for NINJ1 lesion formation. In the plasma membrane of living cells NINJ1 forms face-to-face dimers that sequester the amphipathic transmembrane helix (TM1, red). During cell death NINJ1 dimers associate laterally to form oligomers, assembling a double filament resembling a closed zipper. Membrane tension causes a separation of individual dimers within the double filament, thus allowing a kinking of TM1. TM1 kinking stabilizes interactions between side-by-side NINJ1 neighbors and increases the space between opposing face-to-face NINJ1 molecules, thus supporting the complete separation of the NINJ1 double filament (zipper opening). Upon separation of the double filament, NINJ1 forms large pore-like lesions to permeabilize the plasma membrane.

Previously, the separation of *trans*-autoinhibited NINJ1 dimers was proposed to trigger NINJ1 oligomerization^13^, however we observe that NINJ1 mutants stabilizing the dimer interface still oligomerize. In addition, MD simulations support the lateral association of dimers to form oligomeric filaments as the path to pore formation. Indeed, for NINJ1 to rupture cells, the dimers and tetramers that exist at resting state must first associate into longer filaments, which membrane tension then separates into lesions. While it remains mechanistically unclear what drives the association of NINJ1 dimers into longer filaments, recent structural data proposed the integration of lipids within the NINJ1 filament^15^ potentially implicating changes in lipid composition. Alternative hypotheses implicate the amount or location of NINJ1, and other biophysical properties of the membrane that could promote dimer aggregation, including membrane fluidity. Multiple inhibitors of NINJ1 oligomerization have been reported, including glycine and muscimol^19,33^, and defining their mechanism of action could provide insight into the mechanism of oligomerization.

Dimer stability is critical to determining the membrane-rupturing properties of Ninjurin filaments. In agreement with previous work^13^, we find that dimer stabilizing mutations in NINJ1 prevent PMR. Moreover, we find that the stability of the NINJ2 dimer likely renders NINJ2 filaments resistant to separation, as mutations predicted to decrease the stability of the NINJ2 dimer interface increase NINJ2-mediated PMR. We further observe that mixed filaments of NINJ1 and NINJ2 permeabilize the plasma membrane more slowly than NINJ1 alone, suggesting that the ability of NINJ2 to stabilize filaments in a closed conformation could regulate the permeabilization capability of NINJ1.

The formation of NINJ1 oligomers coincides with the metabolic death and swelling of dying cells. While live cells dynamically react to volume increases, quickly adjusting cell volume and thereby reducing the swelling-associated increase in membrane tension, this is not possible in dying cells. Therefore, in dying cells, swelling leads to a permanent increase in membrane tension, once the pool of free lipids is exhausted. Our AFM measurements of increased membrane tension during ferroptosis coupled with the observation that this tension never builds up when swelling is blocked highlight the key role tension plays in opening closed NINJ1 lesions. While measurements with fluorescent probes, such as Flipper-TR^34^, are another common way of measuring membrane tension, we found that membrane alternations characteristic of ferroptosis invalidated these measurements. The critical role for membrane tension in NINJ1 lesion opening is corroborated by our MD simulations, which confirm the need for membrane tension to separate the hydrophilic interface. The computing requirements of MD simulations mean that simulations must occur in microseconds, rather than the hours it often takes cells to die. To accommodate this compressed timescale, we apply a lateral tension that is higher than the tension measured in our AFM tether pulling experiments, but within the range used in MD simulations of other membrane proteins^27,28^.

Our data, including the visualization of NINJ1 lesions in liposomes and MD simulations, support a double filament zipper model in which NINJ1 causes PMR while remaining associated with the plasma membrane of dead cells^14^, forming a protective protein layer around the lesion. While we cannot exclude that multiple pathways can lead to lesion formation or cell type differences, none of our simulations or experimental results are consistent with the release of NINJ1 from dying cells. Going forward, it will be important to invest effort into understanding what regulates the lateral association of NINJ1 dimers and thus governs filament formation during cell death. Finally, the importance of NINJ1-mediated PMR during health and disease are in their infancy^32,35^ and extensive work is required to understand the potential therapeutic benefits of targeting NINJ1.

## Acknowledgements

EH is supported by EMBO postdoctoral fellowship ALTF 27-2022 and a Novartis Young Investigator Grant. PB is supported by grants from the Swiss National Science Foundation (310030B_198005 and 310030_219286). DM is supported by a grant from the Swiss National Science Foundation through the NCCR Molecular Systems Engineering. VP is supported by grant ANR-22-CE15-0032-01. CS acknowledges support by the Helmholtz Association (VH-NG-1526). DF is supported by the University of Bern. CP is supported by grants from the Swiss National Science Foundation (310030_207974) and NIH grant R35GM158187. SH is supported by the Swiss Nanoscience Institute. KP is supported by the Deutsche Forschungsgemeinschaft under Germany’s Excellence Strategy – EXC 2075 – 390740016 and by the Stuttgart Center for Simulation Science (SC SimTech). KP gratefully acknowledges the scientific support and HPC resources provided by the Erlangen National High Performance Computing Center (NHR@FAU) of the Friedrich-Alexander-Universität Erlangen-Nürnberg (FAU) under the NHR project MoTrNanoMat. NHR funding is provided by federal and Bavarian state authorities. NHR@FAU hardware is partially funded by the German Research Foundation (DFG) – 440719683.

We wish to thank the staff of the Cellular Imaging Facility and Animal Facility at the University of Lausanne for their crucial work maintaining microscopes and managing mouse colonies and all members of the Broz lab for helpful discussions and feedback. We thank Dr. Vishva Dixit (Genentech Inc.) for sharing the anti-NINJ1 antibody, Aaron Ponti (Single cell facility, D-BSSE) for his image processing assistance and Siewert-Jan Marrink and Paulo Cesar Telles de Souza for sharing a development beta version of Martini 3 with us.

## Author contributions

Conceptualization: EH, EMB, KP, DJM, PB

Investigation: EH, EMB, GA, SM, KP, HBL, SAM, MD, MA, JCS, IMG, GC, MS, JH

Data analysis and visualization: EH, EMB, GA, KP, SM, CS, JCS, MD, JH, DJM, PB Writing - first draft: EH, EMB, PB

Writing - revisions: EH, EMB, PB, KP, GA, SAM, DJM, SH

Supervision and funding acquisition: PB, EH, VP, DJM, CS, KP, SH, DF, CP

The authors agree that EH and EMB can list their own names first when referring to this manuscript.

## Data and material availability

All data are available in the main and supplemental figures. All plasmids will be made available on Addgene. Any requests for materials should be made to PB.

## Materials and methods

### Cell culture

BMDMs were differentiated from bone marrow isolated from C57BL/6J WT and *Ninj1^−/−^* mice as previously reported, using L929 cell supernatant as a source of M-CSF. BMDMs were maintained in DMEM high glucose (Thermo Fisher Scientific) supplemented with 10% FCS (BioConcept), 20% MCSF, 1% NEAA (Thermo Fisher Scientific), 10 mM HEPES (Thermo Fisher Scientific), and 1% Pen/Strep (BioConcept) and stimulated on days 9 to 10 of differentiation. WT and *Ninj1^−/−^* iBMDMs were previously reported^11^ and maintained in DMEM supplemented with 10% FCS, 10% MCSF, 1% NEAA and 10 mM HEPES. HeLa and HEK293T cells were maintained in DMEM supplemented with 10% FCS. MOLT-4 cells were maintained in RPMI with 20% FCS. HEK293T-DmrB-Caspase1 cells were previously described^36^. All cells were maintained at 37°C, 5% CO2 in humidified incubators. All cell lines are regularly tested in the lab for mycoplasma contamination and are mycoplasma free.

### Animals

All experiments involving animals were performed under the guidelines of and with approval from the cantonal veterinary office of the canton of Vaud (Switzerland), license number VD3257. All mice were bred and housed at a specific pathogen free facility at 22±1°C room temperature, 55±10% humidity and a day/night cycle of 12h/12h at the University of Lausanne. *Ninj1^−/−^*mice were described previously^14^.

### Plasmids

All cloning for plasmids in this manuscript was done with Infusion cloning (Takara), following the manufacturer’s instructions. All plasmids from this manuscript will be available on Addgene. NINJ1-GFP (Addgene 208779) was previously published^14^. Plasmids used in this study: pMT HA^TMD^-mCherry, pMT HA^TMD^-GFP, pLVX NINJ2-GFP, pLVX NINJ1-mCherry, pLVX NINJ2-mCherry, pLVX Chimera 1 - GFP, pLVX Chimera 2 - GFP, pLVX Chimera 3 - GFP, pLVX Chimera 4 - GFP, pLVX NINJ2^L24V,T29S^ - GFP, pLVX NINJ2^T29N^- GFP, pLVX NINJ2^T29S^ - GFP, Chimera 5 - GFP, pLVX NINJ1^A48S^ – GFP, pLVX NINJ1^L52N^ – GFP, pLVX NINJ2^K31R,S32A^ – GFP, pLVX NINJ1-mEOS, pCDNA-NINJ1, pCDNA-NINJ2, pCDNA NINJ2^K31R,S32A^.

### CRISPR editing

*NINJ1/2* DKO HeLa cells were made by lentiviral transduction of HeLa cells with Cas9 and a sgRNA to *NINJ1* (sequence: GAGGAGTACGAGCTCAACGG) on a blasticidin resistance cassette (pLentiCRISPR Blast, gift of A. Jourdain, University of Lausanne) and then secondarily transducing with lentivirus harboring a *NINJ2* targeting sgRNA (sequence: CATGGCGTTGGACATGAACA) and a zeocin resistance cassette. Lentivirus was made by seeding 1.5 × 10^6^ HEK 293T cells the day before in a 6 well dish. The following day 250 ng pVSV-G, 1250 ng psPAX2 and 1250 ng of the sgRNA-containing vector were transfected with 8.25 μL of LT-1 (Mirus). Media was changed the next morning to 4 mL DMEM supplemented with 10% FBS and 1% BSA (Sigma Aldrich). Lentivirus containing supernatants were harvested 48 h post transfection and filtered through a 0.22 μm syringe filter. Recipient cells were spinfected at 500 × *g* for 90 min with 1 mL (Cas9 containing) or 200 μL (sgRNA only containing vector) of virus-containing supernatant in the presence of 8 μg/mL polybrene (Sigma Aldrich). HeLa cells were selected with 5 μg/mL blasticidin (Invivogen) and 200 μg/mL zeocin (Invivogen).

### Cell stimulations

All cells were seeded the day before stimulation and, unless indicated, stimulations were performed in OptiMEM (Thermo Fisher Scientific). Unless specifically indicated, ferroptosis was induced with 750 μM cumene hydroperoxide (CuOOH, Sigma Aldrich) or 5 μM ML162 (Sigma Aldrich). Pyroptosis was induced with 5 μM nigericin (Sigma Aldrich) for 2 h. Apoptosis was triggered with 500 nM ABT-737 (Lubioscience) + 500 nM S63845 (Lubioscience) for 6 h.

OptiMEM was supplemented with sucrose (Sigma Aldrich), PEG600 (Fuka) or PEG6000 (Santa Cruz) as indicated. Hypotonic media was made by mixing microbiology grade H2O (BioConcept) with OptiMEM. 76 mOsm solution was obtained by mixing 80% H2O with 20% OptiMEM. Before incubation, media was removed from cells, they were washed once with 76 mOsm solution before proceeding with the experiment. Osmolarity of all solutions was measured on a vapor pressure osmometer (Wescor).

### Western blotting

For crosslinking experiments stimulations were either performed directly in PBS containing Ca^2+^ and Mg^2+^ (Thermo Fisher Scientific) or, after stimulation media was changed to PBS with Ca^2+^ and Mg^2+^ to perform crosslinking. At the endpoint 25 mM bis(sulfosuccinimidyl)suberate (BS^3^, Thermo Fisher Scientific) dissolved in water was spiked into the well to a final concentration of 3 mM and cells were incubated at room temperature for 5 min. The reaction was quenched through the addition of an equal volume of 40 mM Tris-HCl pH7.4 and incubated for 15 min at room temperature. Supernatant was then removed and cells lysed in sample buffer (NuPage LDS sample buffer (Thermo Fisher Scientific) + 66 mM Tris-HCl pH7.4, 2% SDS (PanReac) + 10 mM DTT (Thermo Fisher Scientific). Samples were boiled at 97°C for 7 minutes.

Proteins were precipitated from the supernatant by chloroform/methanol extraction. Supernatants were mixed with 1 volume of methanol (Thermo Fisher Scientific) and 0.3 volumes chloroform (Sigma Aldrich), vortexed then centrifuged at 14,000 × *g* for 10 min at 4°C. The upper phase was discarded and 1.3 volumes MeOH added before repeating the vortex and centrifugation steps. The supernatant was discarded and the pellet air dried before resuspending in the corresponding cell lysate or sample buffer.

Samples were then separated on 4-12% BisTris SDS-PAGE gels (Merck) and transferred to nitrocellulose membranes (Amersham GE) using a Transblot Turbo (Bio-Rad). Membranes were blocked in 5% skimmed milk, all antibodies were diluted in 5% milk and incubated either overnight at 4°C or for 1 h at room temperature.

#### Antibodies used

NINJ1 (Genentech Inc, clone 25, 1:8000), H3 (Cell Signaling Technologies 3638S 1:2000), Tubulin-HRP (Abcam ab40742, 1:5000), GFP (Enzo Life Sciences ALX-210-199-R100, 1:1000), mCherry (Abcam ab125096, 1:1000), anti-rabbit IgG HRP (SouthernBiotech 4030-05, 1:5000), anti-mouse IgG HRP (SouthernBiotech 1034-05, 1:5000), NINJ2 (produced commercially by Yenzyn).

#### Antibody generation

Rabbit anti human NINJ2 was made by YenZym antibodies, LLC to a synthetic NINJ2 peptide comprising the first 60 aa of human NINJ2 conjugated to KLH (GenScript).

### Live cell imaging

Live cell imaging was performed on a Zeiss LSM800 equipped with heating and CO2 control using a 63X 1.4 NA oil immersion objective with autofocus (Zeiss Definite Focus.2 system) and data were acquired using Zeiss ZEN 2 software.

BMDMs were seeded at 75,000 cells per well of iBidi 8 well slides the day before stimulation. Cells were stained with CellBrite Orange (Biotium, 1:200 in OptiMEM) for 20 minutes, washed three times in warm OptiMEM and then stimulated as indicated before immediately placing them on the microscope. DRAQ7 (Biolegend) was present at 300 nM. Cells were imaged every 3 min with 10-12 Z planes per image. At least 6 fields of view were imaged per experiment and 3 independent experiments performed.

HeLa cells were seeded at 10,000 cells per well in the afternoon, transfected the following morning using XtremeGene9 (Sigma Aldrich), then NINJ1 expression was induced with 2 µg/mL doxycycline (Sigma Aldrich) in the afternoon before imaging the following day in OptiMEM in the presence of 300 nM DRAQ7 and 750 μM cumene. Cells were imaged every 3 min with 4-6 Z planes per image. 17-29 cells from 2-3 biological replicates were quantified in each condition.

For imaging on the Incucyte SX1 (Sartorius) BMDMs were seeded at 50,000 cells per well of 96 well plates and stimulated as indicated. Brightfield images were acquired using a 10X objective.

### Tether extrusion by atomic force spectroscopy (AFM)

Experiments were performed in BMDM media without the addition of MCSF or FCS. For tether extrusion, we employed an AFM (CellHesion, JPK Instruments) mounted on an inverted optical microscope (Observer.Z, Zeiss). The day before experiments, 200,000 BMDMs were seeded in tissue culture dishes with a glass surface (FluoroDish, WPI). To maintain culture conditions (37°C and 5% CO2), the experimental setup was placed in a temperature-controlled chamber (The Cube, Life Imaging Services) and the dish was continuously perfused with a humidified gas mixture based on synthetic air containing 5% CO_2_ ^37^.

Rectangular OBL-10 cantilevers (Bruker) with V-shaped tips were coated with 2.5 mg/mL Concanavalin A (Sigma) in PBS (phosphate buffered saline) solution (Gibco) for 1 h at 37°C. The cantilevers had nominal spring constants ranging between 0.006 and 0.002 N/m. Before each experiment cantilevers were calibrated using the thermal noise method^38^.

To mechanically extract membrane tethers from cell membranes, single BMDMs were indented by the tip of the microcantilever with an approach speed of 10 µm/s until reaching a setpoint contact force of 0.25 nN. After the contact time between tip and cell of 0.5 s, the tip was retracted at 10 µm/s. During approach and retraction, a force-distance curve was recorded from which the tether force was extracted. The tether force was extracted from single force-distance curves using a custom Python script (https://doi.org/10.5281/zenodo.17177629). For each cell the experiment was repeated three times to record three force-distance curves from which the mean tether force was determined. Membrane tension (Tm) was computed from the tether force (Ft) according to Eq. (1), as described previously ^22–24^. Where B is the membrane bending stiffness. For macrophages we used B = 8×10^-19^ Nm^39^.

### Image analysis

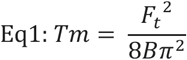

All optical microscopy datasets were analyzed and processed with Fiji software.

The pattern of fluorescent protein-tagged constructs over time on the plasma membrane was evaluated using QuASIMoDOH^40^ (Quantitative Analysis of the Spatial-distributions in Images using Mosaic segmentation and Dual parameter Optimization in Histograms) using the basal plane of cells. The distribution inhomogeneity at each time point (dt) was normalized to the distribution inhomogeneity at the initial time point of the experiment (ti). Time 0 was defined as described below.

To determine the DRAQ7 ratio a maximum projection of all imaged slices was made. An ROI was manually drawn around the nucleus and the mean pixel intensity determined within this ROI at all timepoints. The ratio was then determined using the formula (*mean t* − *mean ti*)/*mean ti* where ti is the value in the first frame imaged. Time 0 in all analyses of NINJ1 inhomogeneity or cell diameter is set as the first frame where this ratio exceeds 1.

To measure the diameter of cells, a straight line was manually drawn across the CellBrite image from one Z plane and the length of this line measured. The diameter was then normalized to the diameter of the cell in the first frame imaged using the formula (*diameter t* − *diameter ti*)/*diameter ti*. For each condition a minimum of 20 cells from 3 biological replicates were analyzed.

### Single-molecule localization microscopy and analysis

For STORM and STORM/PALM imaging, HeLa cells were seeded on 25 mm glass slides. After blocking, cells were incubated with AlexaFluor 647-conjugated anti-GFP nanobodies (FluoTax-X4, NanoTag; 1:500 dilution) in blocking buffer for 2 h. The sample was then washed three times for 10 min in PBS. For dual color STORM/PALM, HeLa cells were transfected with NINJ2-GFP and NINJ1-mEOS.

A Nikon Ti Eclipse NSTORM microscope equipped with a 642 nm laser (F-04306-113, MPB Communications Inc.) was used for STORM imaging to switch molecules to the off state. A 405 nm laser (Cube, Coherent) was used to control the return rate of the fluorophores to the emitting state. A F73-888 quad-band dichroic (Chroma) was used to reflect the laser light on the sample. Emitted light from the sample was collected by the objective lens (Apo TIRF 100x, NA 1.49 Oil, Nikon), passed through the dichroic mirror onto an EMCCD camera (iXon DU-897D, Andor). For additional PALM imaging (Dual-color PALM STORM), we used a 405 nm (Coherent) activation laser to control the photoswitching rate of expressed molecules and a 561 nm (Sapphire, Coherent) laser to excite the photoconverted fraction. Emitted light was directed onto the camera using a dichroic mirror and filtered through a Bright Line HC 609/64 emission filter. The width of a square camera pixel corresponds to 160 nm on the sample. Imaging was performed using an established photo-switching buffer made of 50 mM Tris with 10 mM NaCl, supplemented with 10 % glucose, 50 mM MEA, 0.5 mg/mL glucose oxidase and 40 µg/mL catalase. All chemicals were purchased from Sigma. Typically, 10,000 frames were recorded per field of view with continuous laser exposure at 30 ms integration time. The microscope was controlled using NIS elements (Nikon), single-molecule localization analysis was performed using Decode^41^. Localizations were processed using custom MatLab (Mathworks) code, images were rendered using Thunderstorm^42^. Localizations were clustered using DBSCAN^43^.

### Cytotoxicity assays

LDH release was measured from the supernatant of cell cultures following the manufacturer’s instructions (Sigma Aldrich, 11644793001). 100% lysis was performed using a final concentration of 0.1% TritonX-100. Each experiment was performed in technical triplicate and for 100% and untreated conditions the average of the technical replicates obtained. The %LDH release was calculated using the formula (*Value sample* − *Value untreated*)/(*Value* 100% − *Value untreated*) ∗ 100.

Propidium iodide (PI) uptake was measured using a Cytation5 (Biotek) plate reader or Incucyte SX1 (Sartorius). Briefly, cells were stimulated in the presence of 12.5 μg/mL PI (ThermoFisher Scientific) and directly placed on the instrument for analysis. For plate reader analysis the total fluorescence intensity was measured using an excitation wavelength of 535 nm and an emission wavelength of 617 nm. For Incucyte analysis images of PI and brightfield were captured using a 10X objective with 3 images per well and the integrated density of PI signal per image calculated using Incucyte software. Data was then normalized to 100% and untreated controls. The percentage of PI uptake was then calculated using (*Value sample* − *Value untreated*)/(*Value* 100% − *Value untreated*) ∗ 100. Each experiment was performed in technical triplicate.

Cellular ATP content was assayed using CellTiter-Glo 2.0 (Promega G9241) according to the manufacturer’s instructions. Luminescence was read using a Spectramax ID5 (Molecular Devices) plate reader. The % cell viability was calculated using the formula (*Value sample*/*Value untreated*) ∗ 100.

### Volume measurement by fluorescence exclusion microscopy

Volume measurements were performed in PDMS culture chambers of controlled height (h = 22 µm) in OptiMEM medium supplemented with dyes conjugated with dextran molecules of defined sizes (Alexa647-10kDa dextran from ThermoFisher Scientific at 10 mg/mL and FITC-500kDa from Sigma Aldrich at 25 mg/mL). Acquisitions were performed under a Leica DMi8 microscope at 37°C with a 10x magnification objective (N.A. 0.3).

PDMS molds were made with classical photolithography, a mixture of PMDS and its crosslinker (1:10, Sylgard 184 from Dow Corning) was poured onto the mold, degazed and left for reticulation at 65°C for at least 2 hours. Chambers were bonded with air plasma (Harrick Plasma). Cells were injected directly after bonding and incubated overnight at 37°C under 5% CO2 atmosphere in high glucose DMEM supplemented with 10% (v/v) fetal bovine serum and 1% penicillin-streptomycin.

Image analysis was performed using MatLab (The MathWorks Inc.) with homemade scripts as described in Nadjar et al. 2024^18^. Briefly, cell segmentation was performed on the largest dye (FITC-500kDa dextran) and propagated to images obtained with smaller dyes. Calibration of the relation between the fluorescence intensity and pixel height was made for each field of view and each time frame^44^. Points of interest were defined by users on the basis of volume trajectories in order to extract volume and time values corresponding to swelling onset and dye entries.

PDMS is a hydrophobic material, in addition the surface-to-volume ratio in the FXm chambers is very large. This implies that hydrophobic molecules such as drugs are highly depleted by the surface of FXm chambers. To solve this issue, chambers with constant passive renewal of medium were used. Second, a large range of CuOOH concentrations was tested, no reaction was observed in the chambers below 20 mM while 1 mM was sufficient to induce ferroptosis outside the chambers.

### Immunoprecipitation: GFP-Trap

3×10^5^ HEK293T cells were seeded the day before in a 6-well plate in duplicate and transfected the following day using XtremeGene9 following the manufacturer’s protocol. 24 h later cells were washed in cold PBS and then pipetted into 1.5 mL of cold PBS. Cells were pelleted at 300 × *g* for 5 min at 4°C and then lysed in 500 mL PBS + 0.1% TritonX-100 + 1 mM MgCl2 + protease inhibitors (Sigma Aldrich) for 1 h on ice. Nuclei were pelleted by spinning at 20,000 × *g* for 15 min at 4°C and then supernatant (cytoplasm) was removed. 25 µL GFP-Trap magnetic agarose beads (ChromoTek) per sample were washed 3x with PBS and then added to the cytoplasmic lysate for 3 h at 4°C rotating. Beads were then separated and washed 3x with the lysis buffer on a magnet, then resuspended in 2X NuPage LDS sample buffer (Thermo Fisher Scientific) and boiled for 5 min at 95°C followed by Western blotting.

### Molecular dynamics simulations

Active-state NINJ1, i.e. with kinked TM1, was based on the PDB: 8SZA structure^15^ and comprised residues N39-K143. Inactive NINJ1, i.e. NINJ1 with straight TM1, was based on the PDB: 9BIA structure^13^ and comprised residues N33-Q142 with Q45 mutated back to K. The structures were energy minimized using 1000 steepest descent energy minimization in vacuo first. The plasma membrane model with symmetric membrane leaflets reflecting the cell-death state consisted of POPC, POPE, POPS, and cholesterol in 60:5:5:30 molar ratio, solvated by water and 0.15 M NaCl.

All simulations were performed by the simulation engine GROMACS version 2023^45^.

#### Coarse-grained MD simulations

NINJ1 was coarse-grained using martinize2^46^ to the development beta version of Martini 3^47^. The interaction matrix can be found under https://github.com/Martini-Force-Field-Initiative/martini-forcefields/tree/main/martini_forcefields/regular/M3_LEGACY. The secondary structure of the CG proteins was stabilized by elastic bonds (RubberBands^48^) with 500 kJ/mol/nm among the backbone beads within 1.2 nm. Notably, when speaking about “allowing TM1 to kink” in the text, then rubberbands corresponding to the 8SZA structure have been applied to the 9BIA-initiated simulation. Then surrounding was generated by the tool *insane*^49^ and the systems were energy minimized in a three-step energy minimization of 5000 steps each. In the first step the protein and in the second step its backbone were frozen. After generation of velocities 5000 steps of equilibration while position restraining the protein were performed with the time step of 10 fs. Afterwards, 20 fs time step was used to perform 10 ns of equilibration while position restraining the protein and 10 ns while position restraining the backbone beads only. In the equilibration steps the semiisotropic Berendsen barostat^50^ with the time constant of 12 ps was used and the position of the center of mass of the reference coordinates was scaled and the compressibility was 3e-5 bar^-1^. In the production run the c-rescale barostat^51^ with the time constant of 10 ps in the semiisotropic mode was used, the compressibility amounted to 3e-4 bar^-1^ and the center of mass of the system was linearly removed every 500 steps. In all CG simulations the temperature was controlled by the v-rescale thermostat^52^ with the time constant of 1 ps for protein/membrane and solvent separately to be at 293 K. The neighbor list was updated every 20 steps and the Verlet cut-off scheme^53^ with buffer tolerance of 0.005 kJ/mol/ps per atom was used. The electrostatic interaction behind 1.1 nm were calculated by reaction-field, the relative dielectric constant amounted to 15, and the van der Waals forces were potential-shifted to be zero at 1.1 nm. In the simulations of 60mer NINJ1 pores the phosphate z-positions were position restrained by 2 kJ/mol/nm potential mimicking the adsorption of the membrane on mica. Membrane tension was generated by applying either 5 or 10 bar along the z-axis. Membrane tension, Γ, was calculated as Γ = *L*z (*P*z - 〈*P*xy〉)^28^ with *L*z being the size of the simulation box in the *z*-direction, here ∼10 nm, *P*z is the pressure along the *z* axis, and *P*xy the average pressure in the *x*,*y* plane. For simplicity we refer to the resulting membrane tension as 5 or 10 mN/m membrane tension.

##### All-atom MD simulations

The all-atom systems were prepared using our multiscaling methodology^54^. In detail, the systems were generated and equilibrated at coarse-grained resolution, following the methodology and settings described in the previous paragraph. Then, *backward*^55^ was used to transform the systems back to all-atom resolution using the CHARMM36m force-field^56,57^ and the TIP4P water model^58^. The backmapped proteins were replaced by the original energy minimized NINJ1 structures and the overlapping water molecules removed. The systems underwent the same 3-steps energy minimization as described for CG simulations above. Then velocities were generated and the protein surroundings was relaxed in a three-step equilibration with position restraints on the proteins (performing 5000 steps with 0.2 fs time step and 1-ns simulation with 2 fs time step) or on the backbone in the latest step (10 ns with 2 fs time step). During those simulations, the Berendsen barostat with semiisotropic pressure coupling scheme, time constant of 5 ps, reference pressure 1 bar and compressibility of 4.5e-5 bar-1 were used and the center of mass scaling of reference coordinates was performed. The temperature of the protein/membrane and water/ion groups were controlled individually to be at 293 K by the Nosé-Hoover thermostat^59^ and a time constant of 0.5 ps. In the production run simulations the same settings were used except for pressure being coupled by the Parrinello-Rahman barostat^60^ with the time constant of 5 ps and the center of mass of the system was linearly removed every 100 steps. In MD simulations of endless filaments, anisotropic pressure coupling scheme was used with diagonal elements equaling to 0. Membrane tension was generated by applying enhanced pressure (50 or 100 bar in AA simulations) along the membrane normal. Due to the initial box size in the z-direction of ∼10 nm, the resulting membrane tension Γ in the membrane plane amounts to 50 or 100 mN/m, respectively.

In all all-atom simulations LINCS^61^ were used to constrain bonds to hydrogen atoms, particle-mesh Ewald^62^ was used to calculate electrostatic interactions behind 1.2 nm, while the Lennard-Jones interactions were potential-switched to zero from 0.8 to 1.2 nm. The neighbor list was updated every 10 steps and the Verlet cut-off scheme with buffer tolerance of 0.005 kJ/mol/ps per atom was utilized.

### Preparation of samples for AFM of liposomes

Purified NINJ1 and proteoliposomes were prepared as previously described^14^. NINJ1 proteoliposomes were diluted 1:45 in buffer solution (150 mM NaCl, 20 mM HEPES, pH 7.5) and adsorbed onto a freshly cleaved mica disc of 6 mm diameter at room temperature. After an incubation time of 45 min, during which the proteoliposomes adsorbed onto the mica and fused into a continuous supported lipid membrane (SLM), the sample was gently rinsed with imaging buffer solution (300 mM NaCl, 40 mM HEPES, pH 7.5). Every experimental condition was reproduced at least three times using freshly prepared proteoliposomes or liposomes (control), new AFM supports and cantilevers.

### AFM imaging

FD-based AFM^63,64^ was performed with a Nanoscope Multimode 8 (Bruker, USA) operated in PeakForce Tapping mode. The AFM was equipped with a 120 μm piezoelectric scanner and fluid cell. The AFM cantilevers used had a nominal spring constant of 0.1 N m^-1^ resonance frequency of ∼ 110 kHz in buffer solution and a sharpened silicon tip with a nominal radius of 8 – 10 nm (BioLever mini BL-AC40, Olympus Corporation). AFM topographs were recorded in imaging buffer (300 mM NaCl, 40 mM HEPES, pH 7.5) at room temperature as described^65^. Briefly, the maximum force applied to image the samples was limited to ∼ 100 pN and the oscillation frequency and oscillation amplitude of the cantilever were set to 2 kHz and 20 nm, respectively. Topographs were analyzed using the AFM analysis software (NanoScope Analysis 1.8). The AFM was placed inside a home-built temperature controlled acoustic isolation box.

### Statistical analysis

All graphs were made and statistical analyses performed using Graphpad Prism v10. Details of all analyses, including number of replicates and tests performed can be found in the figure legends. In all cases, a p value of less than 0.05 was used to determine statistical significance.

## Supplementary Figures

**Figure S1.**
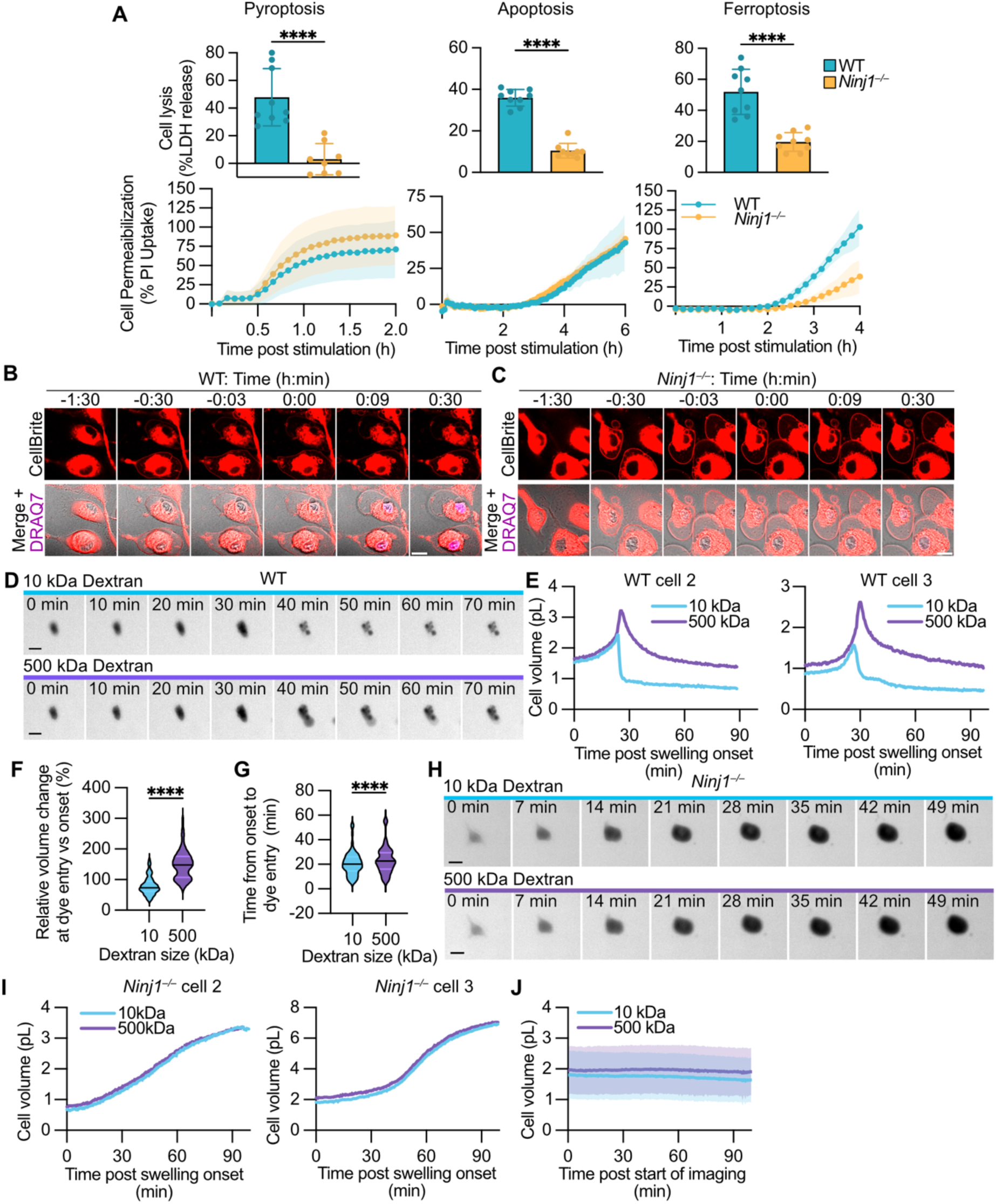
Ferroptotic cell death is characterized by NINJ1-dependent DAMP release and swelling. **(A)** WT and *Ninj1^−/−^* BMDMs were treated with nigericin (pyroptosis, 2h), ML162 (ferroptosis, 4h) or ABT737 and S63845 (apoptosis, 6h) and analyzed for LDH release and PI uptake. (**B-C**) Stills of CellBrite from live cell imaging of WT (B) and *Ninj1^−/−^* (C) BMDMs treated with ML162 in the presence of DRAQ7. Scale bar 10 μm. Corresponds to Figure 1A&C. **(D)** Stills from FXm of WT iBMDMs treated with cumene. Scale bar 10 μm. **(E)** Volume measurements of 2 representative WT iBMDMs treated with cumene using FXm in the presence of 10 kDa Alexa647 Dextran and 500 kDa FITC Dextran. Time 0 indicates the onset of swelling. **(F)** Quantification of the relative change in cell volume of WT iBMDMs treated with cumene from onset of swelling to the uptake of 10 kDa and 500 kDa Dextrans. **(G)** Quantification of the time between the onset of swelling and uptake of 10 kDa and 500 kDa Dextrans by WT iBMDMs treated with cumene as measured by FXm. **(H)** Stills from FXm of *Ninj1^−/−^* iBMDMs treated with cumene. Scale bar 10 μm. **(I)** Volume measurements of two representative *Ninj1^−/−^* iBMDMs treated with cumene using FXm in the presence of 10 kDa Alexa647 Dextran and large 500 kDa FITC Dextran. Time 0 indicates the onset of swelling. **(J)** Volume measurements using FXm of WT iBMDMs treated with ethanol in the presence of 10 kDa Alexa647 Dextran and 500 kDa FITC Dextran. Time 0 indicates the start of imaging. Data in A are mean ± SD from 2-3 independent experiments and were analyzed by unpaired t test with Welch’s correction. Data in (B&C) are representative of 3 independent experiments. Data in (D&H) are representative of 2 independent experiments. Each graph in (E&I) are values from one cell and are representative of at least 34 cells. Data in (F&G) are from 60 cells from 2 independent experiments and were analyzed by two-tailed Wilcoxon matched-pairs signed rank test. **** p<0.0001. Data in (J) are mean ± SD from 24 cells from 1 experiment.

**Figure S2.**
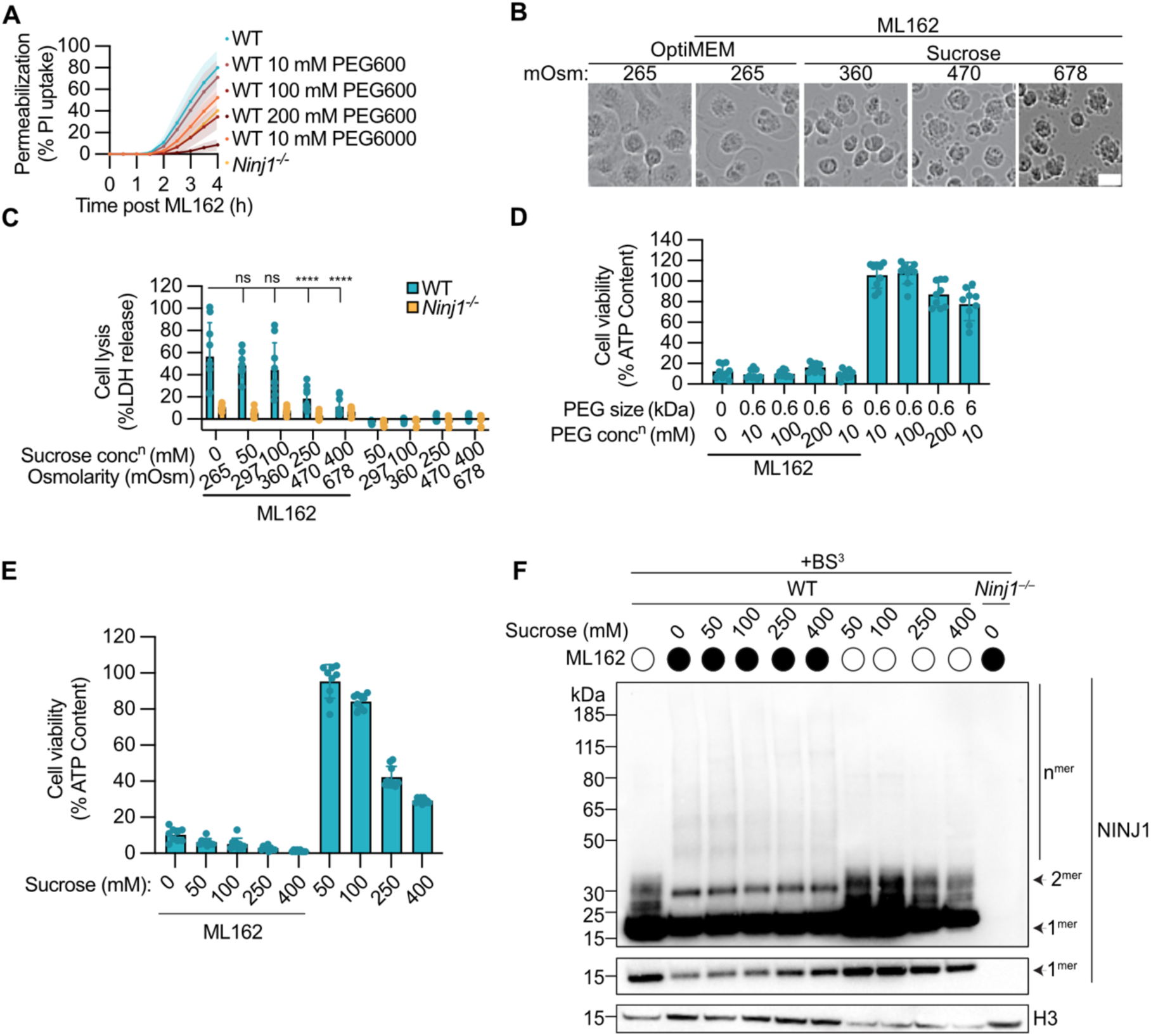
Osmolytes block necrotic swelling and inhibit NINJ1-dependent DAMP release but not NIINJ1 oligomerization. **(A)** PI uptake by WT and *Ninj1^−/−^* BMDMs treated with ML162 in media containing the indicated concentrations of PEG600 or PEG6000. **(B)** Phase contrast stills from Incucyte of WT BMDMs treated with ML162 in the presence of the indicated osmolarities of sucrose for 4 h. Scale bar 20 μm. **(C)** LDH release from WT and *Ninj1^−/−^* BMDMs treated with ML162 for 4 h in the presence of the indicated concentrations of sucrose. **(D&E)** Viability of WT BMDMs treated with ML162 in the presence of the indicated concentrations of PEG600, PEG6000, or sucrose for 4 h by measuring intracellular ATP. **(F)** Crosslinking immunoblot of NINJ1 oligomerization in WT and *Ninj1^−/−^* BMDMs treated with ML162 for 4 h in the presence of the indicated concentrations of sucrose. Data in (A, C & D) are mean ± SD from 3 independent experiments, analyzed by 2-way ANOVA followed by Šídák’s multiple comparisons test. Data in (B) and (E) are representative of 3 independent experiments. ns not significant, **** p<0.0001.

**Figure S3.**
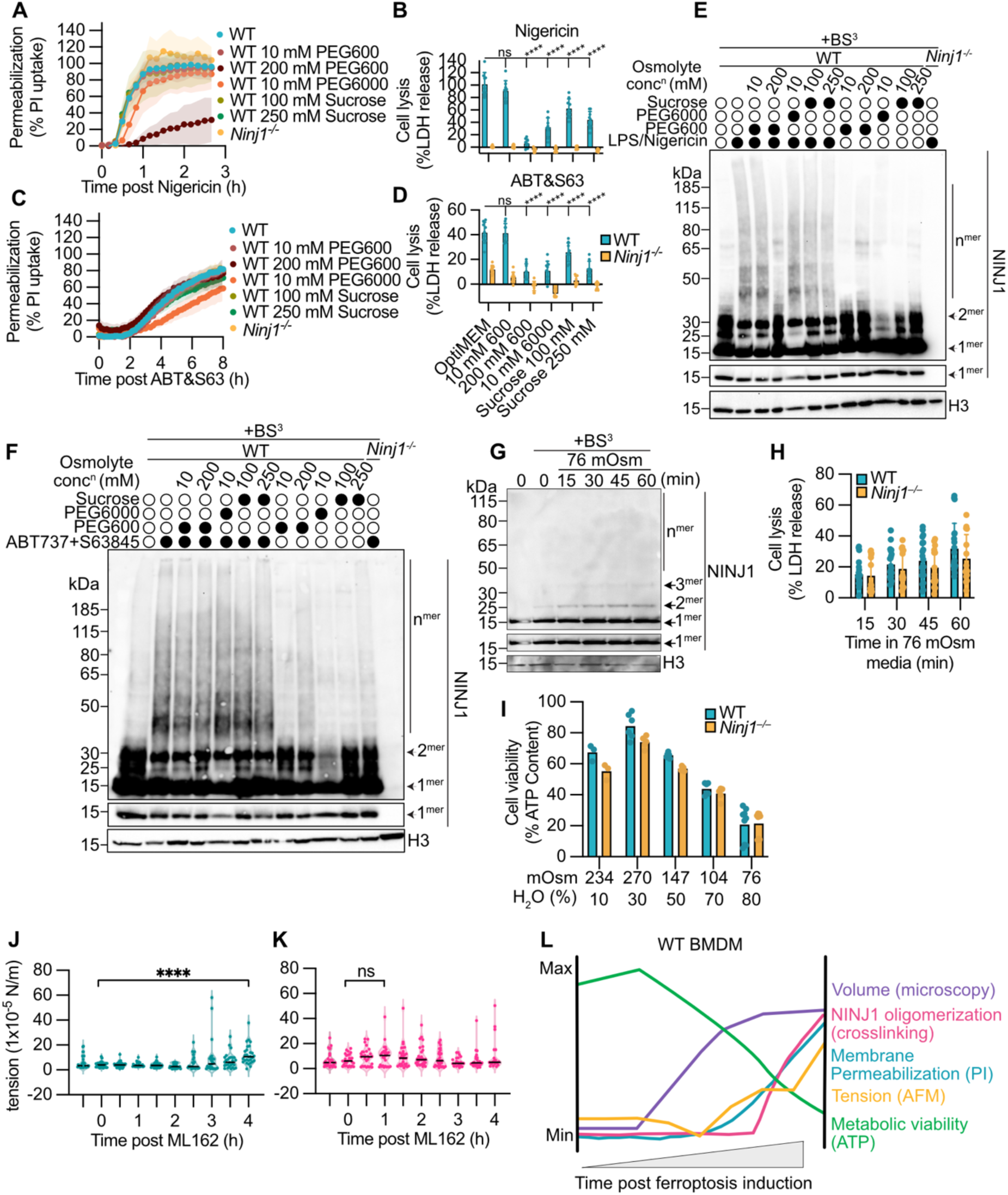
Osmolytes block NINJ1-mediated PMR during pyroptosis and apoptosis. **(A-D)** PI uptake (A,C) and LDH release (B,D) from WT and *Ninj1^−/−^* BMDMs treated with nigericin (A,B) or ABT737+S63845 (C,D) in the presence of the indicated concentrations of PEG and sucrose. LDH release was performed at 2 h for nigericin and 6 h for ABT737+S63845 treatment. **(E-F)** Crosslinking immunoblots from BMDMs treated with nigericin (E, 2 h) or ABT+S63945 (F, 6 h). **(G-H)** Crosslinking immunoblot of NINJ1 oligomerization (G) and LDH release (H) from WT BMDMs incubated with 76 mOsm media for the indicated times. **(I)** Viability of WT and *Ninj1^−/−^* BMDMs treated with the indicated osmolarities of media for 30 min determined by measuring intracellular ATP. **(J-K)** Plasma membrane tension in ML162 treated WT BMDMs in the absence (F) or presence (G) of 10 mM PEG6000. All compounds were added at time 0. **(L)** Schematic overlay of the timing of the labeled parameters during cell death of WT BMDMs, based on data acquired as indicated. Data in (A-D) and are mean ± SD from 3 independent experiments, analyzed by 2-way ANOVA followed by Šídák’s multiple comparisons tests, ns not significant, **** p<0.0001. Data in (E-G) are representative of 3 independent experiments. Data in (H) are mean ± SD from 6 independent experiments. Data in (I) are mean ± SD from 3 independent experiments. Data in J and K are from 3 independent experiments, analyzed by 2-way ANOVA followed by Dunnet’s multiple comparisons test.

**Figure S4.**
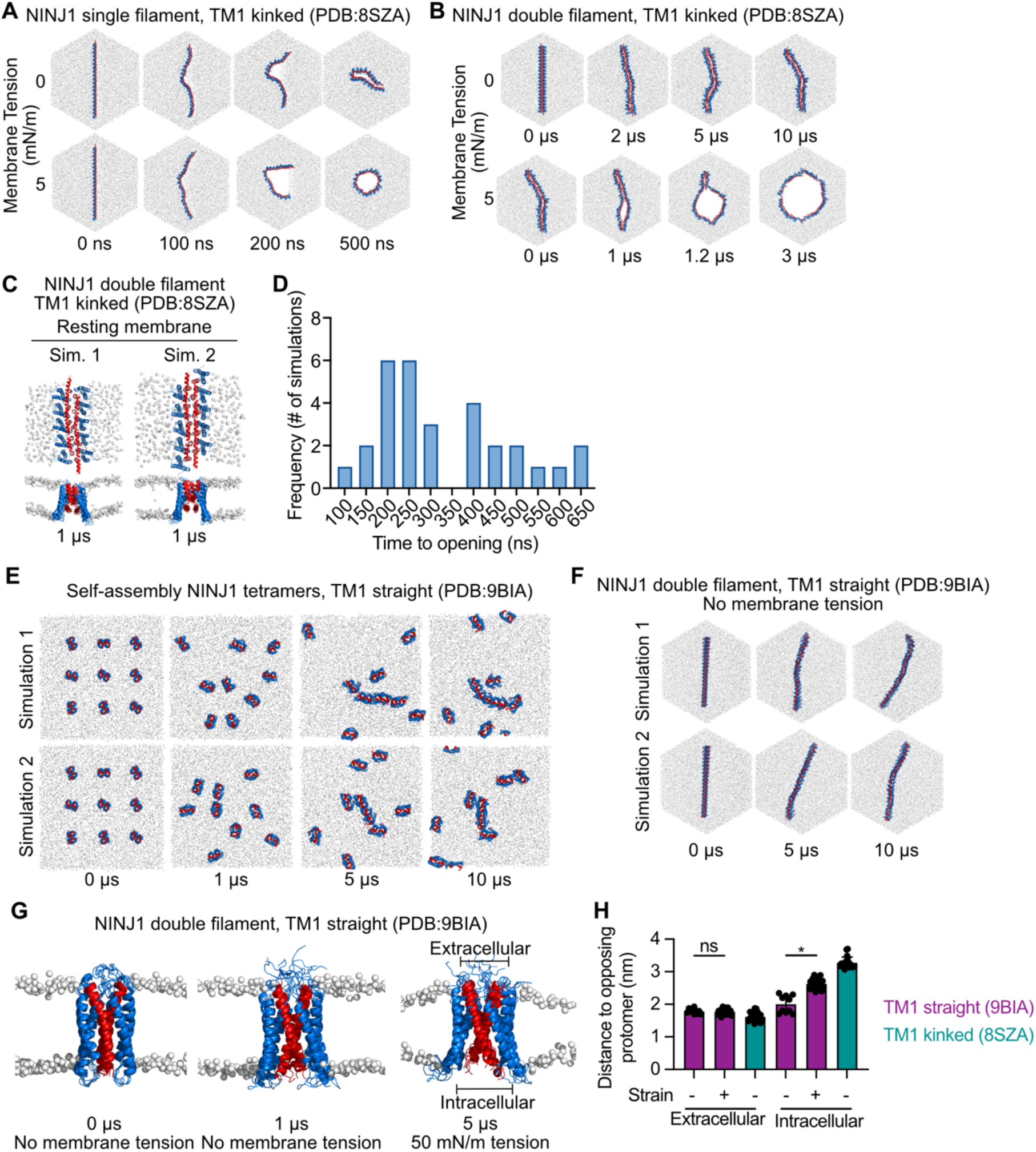
CG and AA MD simulations showing opening of NINJ1 double filaments under membrane tension. **(A)** Stills from 2 representative CG MD simulations of a 20mer NINJ1 single filament (PDB:8SZA) in a membrane in the absence (n=8) or presence (n=5) of 5 mN/m membrane tension. Individual hexagons show snapshots at the indicated times of the simulation. (Expands main Figure 3A). **(B)** Stills from CG MD simulations of a 20mer NINJ1 (PDB:8SZA) double filament arranged with their hydrophilic faces towards each other in the absence (n=3) or presence (n=3) of 5 mN/m membrane tension. (Expands main Figure 3B). **(C)** Stills from 2 representative all-atom MD simulation of an endless NINJ1 (PDB: 8SZA) filament in the absence of membrane tension (n=4). Each protofilament contains 5 NINJ1. **(D)** Frequency distribution showing the time of opening over all all-atom MD simulations of an endless NINJ1 (PDB: 8SZA) double filament under 100 mN/m membrane tension (Related to stills in Figure 3C). **(E)** Stills from 2 representative CG MD simulations of NINJ1 (PDB: 9BIA) tetramers with snapshots at the indicated times of the simulation (n=3). **(F)** Stills from 2 representative CG MD simulations of a 20mer NINJ1 (PDB:9BIA) double filament with a straight TM1 in the absence of membrane tension (n=3). **(G)** Stills of all-atom MD simulations of endless NINJ1 (PDB:9BIA) double filaments with straight TM1 in the absence (n=2) and in the presence (n=5) of 100 mN/m membrane tension. Each protofilament contains 5 NINJ1. Lines on the right correspond to quantification in (H). **(H)** Spacing of the helical bundle of each opposing protomer (n=5 NINJ1 pairs per simulation) at the extracellular and intracellular end of the endless NINJ1 double filament with either straight TM1 (visualized in G, estimated in simulations in absence or presence of membrane tension) or with kinked TM1 (visualized in B, estimated in simulations without tension). In all MD simulations red represents TM1 and blue the remaining structured part of the protein, the lipid phosphates are shown as white spheres. Data in (H) are mean ± SD of measurements from 10-25 NINJ1 pairs, analyzed by one-way Kruskal-Wallis ANOVA and Dunn’s multiple comparisons test, ns not significant, * p< 0.05.

**Figure S5.**
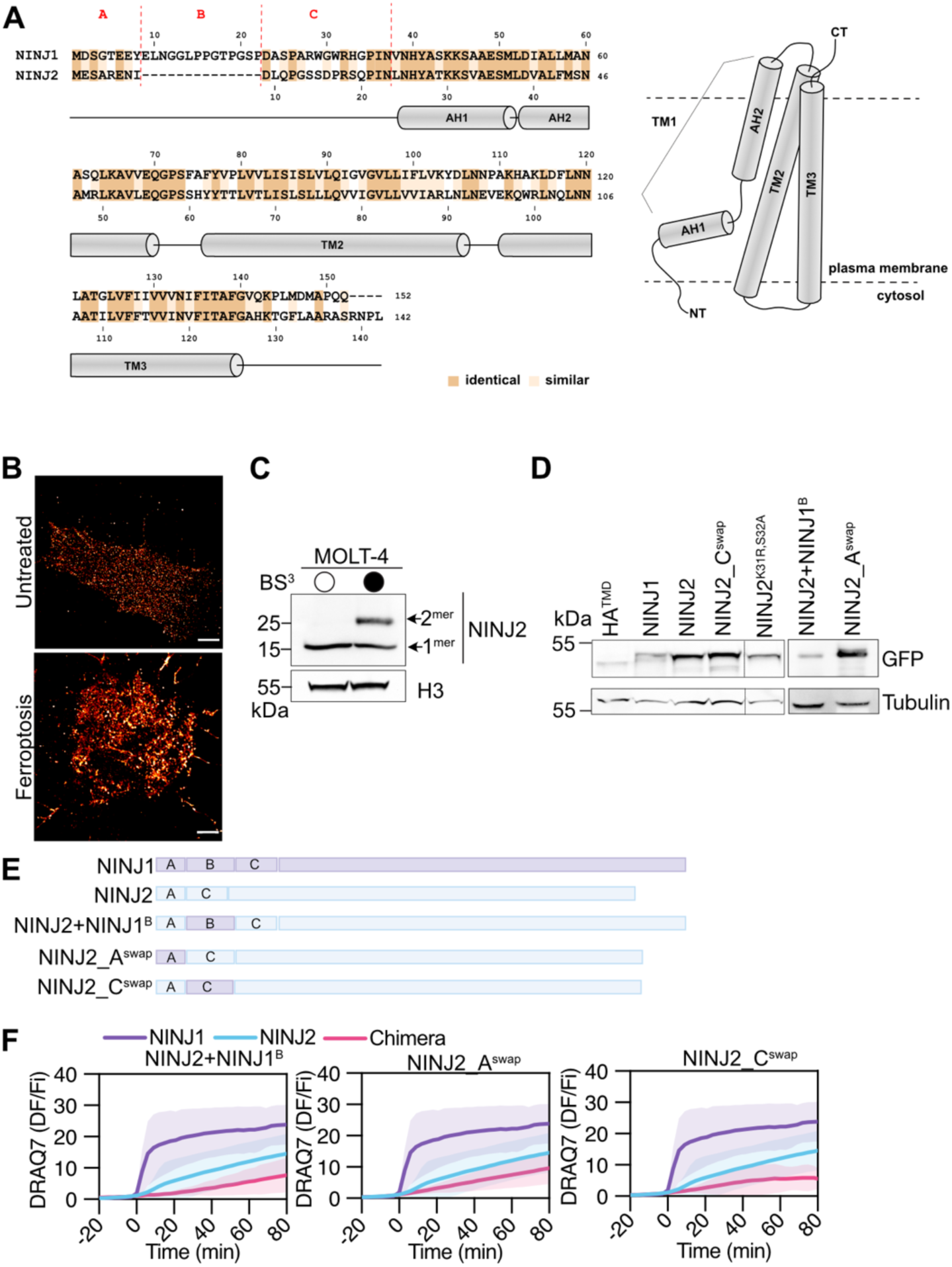
Functional Characterization and Site-Directed Mutagenesis of NINJ2. **(A)** (Left) Alignment of human NINJ1 and NINJ2. The sequences corresponding to transmembrane (TM) helix 1, 2 and 3 are shown and the kink in TM1 separates it into alpha helix (AH)1 and AH2. (Right) A model of NINJ1 in the plasma membrane of cells showing kinked TM1. **(B)** Representative STORM images of untreated or Cumene (ferroptosis) treated *NINJ1/2* dKO HeLa cells expressing NINJ2-GFP. Scale bar 2 μm. **(C)** Immunoblot of expression of NINJ2 and H3 in human MOLT-4 cells ± the crosslinking agent BS^3^. **(D)** Expression of NINJ1, NINJ2, mutant and control constructs tagged with GFP in HEK293Ts. Tubulin serves as a loading control. **(E)** Schematic of NINJ1 and NINJ2 WT highlighting the corresponding regions mutated in each chimera. Regions A, B, and C correspond to the sequences indicated in (A). **(F)** DRAQ7 ratio measured over time from maximum intensity projections of the indicated constructs from live cell imaging performed in *NINJ1/2* dKO HeLa cells treated with cumene in the presence of DRAQ7. Time 0 indicates the first detectable DRAQ7 uptake. The same traces for NINJ1 and NINJ2 are shown with each chimera or mutant. Data in (B) are representative of 2 independent experiments. Data in (F) are from 3 independent experiments with 16-30 cells each. Lines show the mean ± SD. (D) is representative of three independent experiments.

**Figure S6.**
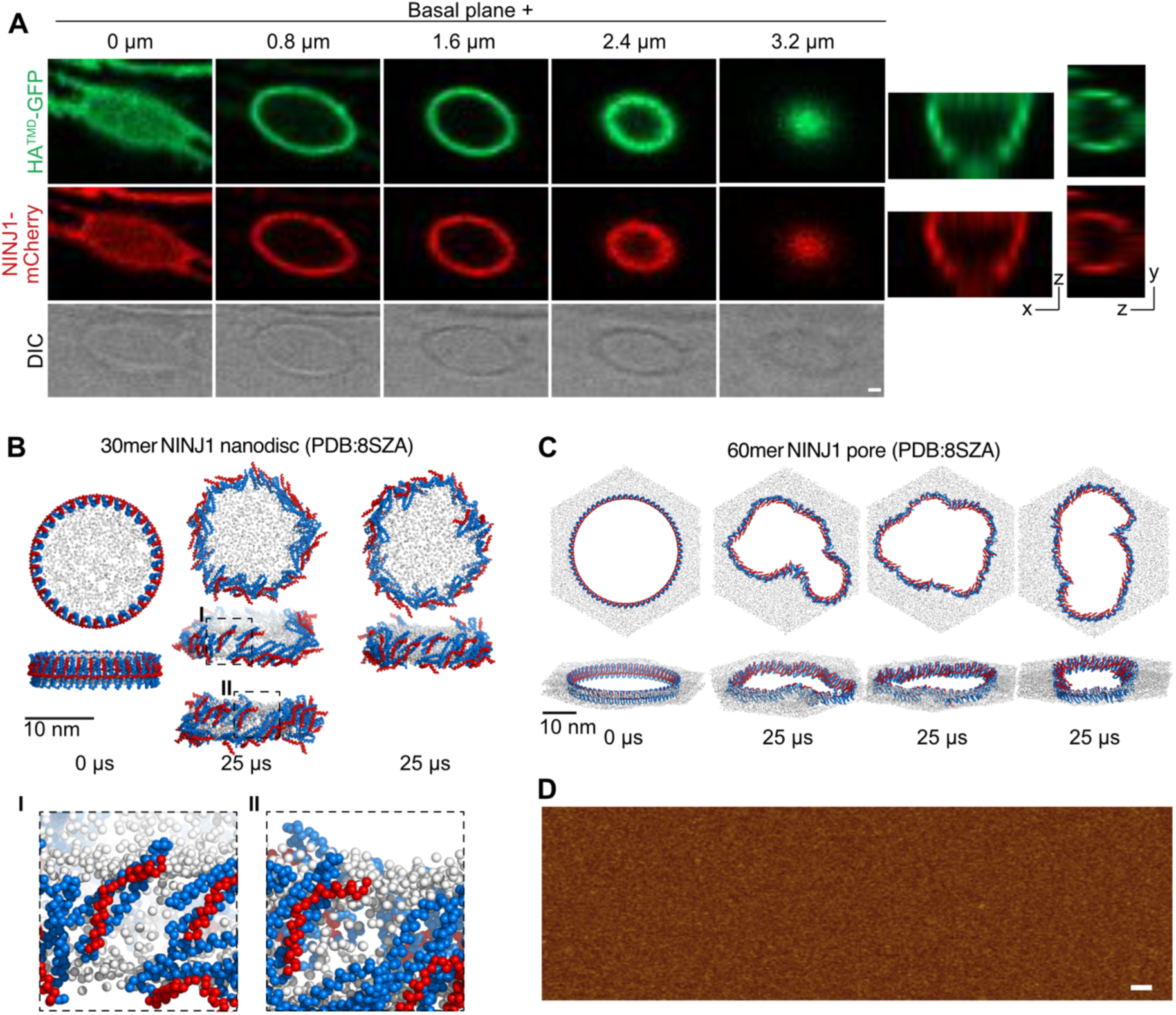
NINJ1 forms membrane pore-like membrane lesions. **(A)** Close up of z-stacks of a vesicle-like structure from live cell imaging of cumene treated *NINJ1/2* dKO HeLa cells expressing NINJ1-mCherry and HA^TMD^-GFP in the presence DRAQ7. Inserts on the right-hand side show a z/x and z/y reconstruction of the structure. Scale bar 1uM. **(B)** CG MD simulations of a lipid nano disc surrounded by a 30mer NINJ1 (PDB:8SZA) single filament (n=6). The same starting point and 25 µs timepoints from two different simulations are shown. Insets I and II show closeups of NINJ1 on the nano disc. **(C)** CG MD simulations of the pore formed by a 60mer NINJ1 (PDB:8SZA) single filament. The same starting point and 25 µs timepoints from 3 different simulations are shown (n=5). Red represents TM1 and blue the remaining structured part of the protein, the lipid phosphates are shown as white spheres. **(D)** AFM topograph of liposomes adsorbed on freshly cleaved mica in the absence of NINJ1. Scale bar, 50 nm. Images in A are representative of 2 independent experiments.

